# Regulation of Eag by ca^2+^/calmodulin controls presynaptic excitability in *Drosophila*

**DOI:** 10.1101/215293

**Authors:** Peter Bronk, Elena A. Kuklin, Srinivas Gorur-Shandilya, Chang Liu, Timothy D. Wiggin, Martha L. Reed, Eve Marder, Leslie C. Griffith

**Affiliations:** Department of Biology, National Center for Behavioral Genomics and Volen Center for Complex Systems, Brandeis University, Waltham, MA, 02454-9110 USA

## Abstract

*Drosophila ether-à-go-go (eag*) is the founding member of a large family of voltage-gated K^+^ channels, the KCNH family, which includes Kv10, 11 and 12, (Ganetzky et al. 1999). Concurrent binding of calcium/calmodulin (Ca^2+^/CaM) to N-and C-terminal sites inhibits mammalian EAG1 channels at sub-micromolar Ca^2+^ concentrations (Schonherr et al. 2000), likely by causing pore constriction (Whicher and MacKinnon 2016). Although the *Drosophila* EAG channel was believed to be Ca^2+^-insensitive (Schonherr et al. 2000), both the N-and C-terminal sites are conserved. Here we show that *Drosophila* EAG is inhibited by high Ca^2+^ concentrations that are only present at plasma membrane Ca^2+^ channel microdomains. To test the role of this regulation *in vivo*, we engineered mutations that block CaM-binding to the major C-terminal site of the endogenous *eag* locus, disrupting Ca^2+^-dependent inhibition. *eag* CaMBD mutants have reduced evoked release from larval motor neuron presynaptic terminals and show decreased Ca^2+^ influx in stimulated adult projection neuron presynaptic terminals, consistent with an increase in K^+^ conductance. These results are predicted by a conductance-based multi-compartment model of the presynaptic terminal in which some fraction of EAG is localized to the Ca^2+^ channel microdomains that control neurotransmitter release. The reduction of release in the larval neuromuscular junction drives a compensatory increase in motor neuron somatic excitability. This misregulation of synaptic and somatic excitability has consequences for systems-level processes and leads to defects in associative memory formation in adults.

**New and Noteworthy:** Regulation of excitability is critical to tuning the nervous system for complex behaviors. We demonstrate here that the EAG family of voltage-gated K^+^ channels exhibit conserved gating by Ca^2+^/CaM. Disruption of this inhibition in *Drosophila* results in decreased evoked neurotransmitter release due to truncated Ca^2+^ influx in presynaptic terminals. In adults, disrupted Ca^2+^ dynamics cripples memory formation. These data demonstrate that the biophysical details of channels have important implications for cell function and behavior.

## Introduction

The repolarization of neurons and muscle cells by K^+^ channels plays an important role in limiting the excitability of cells and returning them to a baseline membrane potential to be ready for the next action potential. K+ channels also play an indirect role in regulating Ca^2+^ levels inside the cell, preventing excitotoxicity and shaping the temporal profile of cellular responses to activity. Several families of Ca^2+^-activated K^+^ channels (e.g. small and large conductance, SK and BK channels), appear to act in a feedback mode, with internal Ca^2+^ activating the channels to hyperpolarize the cell and limit additional Ca^2+^ entry. However, there is one Ca^2+^-regulated K^+^ channel family that appears to act in a counterintuitive manner. The mammalian Ether-à-go-go (EAG) K^+^ channel is completely inhibited at all membrane voltages in the presence of 100-300 nM Ca^2+^/calmodulin (Ca^2+^/CaM; Schonherr et al. 2000; Stansfeld et al. 1996; Whicher and MacKinnon 2016; Ziechner et al. 2006). We sought to understand the role of this seemingly maladaptive regulation by disrupting the CaM interaction with EAG in *Drosophila* where the presence of only one gene coding for EAG simplifies the analysis.

The *Drosophila ether-à-go-go* (*eag*) gene encodes a voltage-gated delayed rectifier K^+^ channel subunit, EAG. The first *eag* mutant was discovered in *Drosophila* and was identified by its ether-induced leg shaking (Kaplan 1969). Subsequently, *Drosophila* EAG was found to define a family of K^+^ channels with multiple homologs in mammals (Warmke and Ganetzky 1994). EAG is a member of the KCNH family of K^+^ channels and is also referred to as Kv10 (Gutman et al. 2003). Although the transmembrane domains of EAG are similar in structure to Shaker-type voltage-gated channels (Warmke and Ganetzky 1994), recent structural data has hypothesized that EAG has an additional novel gating mechanism. This alternate gating mechanism allows cytoplasmic factors like CaM to act on the pore region through the domain linking the C-terminus to the S6 transmembrane domain (Whicher and MacKinnon 2016). Channel closure is achieved by the simultaneous binding of Ca^2+^/CaM to both an N-terminal and a C-terminal site. This requirement for binding at two sites means that disruption of a single CaM binding domain (CaMBD) should block Ca^2+^-dependent gating entirely.

The biophysical properties of EAG have been determined in heterologous expression systems, and little is known about its function in the nervous system. Fly loss-of-function *eag* mutants, in which the conductance is completely missing, have learning deficits (Griffith et al. 1994b) in addition to a robust hyperexcitability in larval motor neurons that also causes spontaneous neuronal firing (Drysdale et al. 1991; Ganetzky and Wu 1983; Griffith et al. 1994b; Srinivasan et al. 2012). In cockroaches, knock-down of EAG revealed a role for Ca^2+^-dependent inhibition in the light response (Immonen et al. 2017). EAG1 knock-out mice have normal learning and memory, sensorimotor function, social behavior, anxiety and only display a mild hyperactivity (Ufartes et al. 2013). However, the existence of multiple EAG family members in mammals raises the possibility of compensation or redundancy and complicates *in vivo* analysis. Cellular physiology in EAG1 knock-out mice has revealed enhanced synaptic facilitation during high frequency stimulation (>= 50 Hz) at the parallel fiber-Purkinje cell synapse in the cerebellum, accompanied by elevated presynaptic Ca^2+^ (Mortensen et al. 2015). In both insects and mammals, the dominant phenotype of loss of EAG channels appears to be increased presynaptic release.

The role of gating by Ca^2+^/CaM *in vivo* is likely to be more complicated, and understanding the role of Ca^2+^ inhibition requires generating an EAG channel that is voltage-gated, but lacks Ca^2+^/CaM inhibition. Such a channel would be predicted to pass more current when Ca^2+^ is high. In this study we show that fly EAG is inhibited by Ca^2+^/CaM, and we generate mutant alleles disrupting the CaM interaction with EAG to interrogate its function. We find that without Ca^2+^/CaM inhibition of EAG, evoked synaptic currents are reduced, and presynaptic Ca^2+^ is reduced during high frequency stimulation. These defects lead to changes in somatic excitability that go in the opposite direction, likely the result of homeostasis. The complex misregulation of excitability in these mutants disrupts higher level behavior, demonstrating that the biophysical details of channel regulation can have profound effects at the organismal level.

## Materials and Methods

### *Drosophila* crosses for recording and protein analysis

Crosses to produce larvae for electrophysiology were set up in vials with 5-6 females and 5 males for each genotype to have consistent numbers of larvae between genotypes and kept at around 22° C in the lab. Crosses were flipped every 3-5 days, but no more than 4 times. Bottle crosses were used for protein extraction. The single alleles analyzed were hemizygous males resulting from a cross of homozygous control or CaMBD mutant females to *w* males. The transheterozygous alleles were females resulting from crossing 2 homozygous alleles of control (WT12 and WT8) or CaMBD mutants (CaMBD8 and CaMBD17). Whenever possible the transheterozygous combinations were used to reduce the effect of potential off target mutations not eliminated during outcrossing of stocks.

### Generation of mutant strains

Control and Calmodulin binding domain (CaMBD) mutant knock-in strains were generated by targeted ends-out homologous recombination using methods described in detail previously (Staber et al. 2011). The homologous arms were amplified by PCR from genomic DNA and each arm was cloned into a TOPO-TA vector (Invitrogen, Thermo Fisher) and were a kind gift from the laboratory of Robert Reenan at Brown University. Mutations to the arm spanning the C2 CaMBD were made by site directed mutagenesis (Stratagene) using primers CGTCCGGAAGATATCCTCCAAATCTCGTCGCACTCCGC (forward) and GCGGAGTGCGACGAGATTTGGAGGATATCTTCCGGACG (reverse). The arms were cloned sequentially into a P[w25.2] target vector. The full *eag* deletion mutant was generated using piggyBack insertions containing FRT sites (Thibault et al. 2004) as described previously (Parks et al. 2004). Briefly, Exelixis stocks PBac{RB}e03618 and PBac{WH}f02960 (p-element on X chromosome) were used to delete the entire *eag* gene. PBac{RB}e03618/FM7a flies were crossed to hs-FLP/Bal and male progeny were crossed to PBac{WH}f02960 females. After 48 hours, the bottle crosses were heat shocked in a 37° C water bath for 1 hour to activate the FLP recombinase. The parents were removed after 72 hours and the bottles were heat shocked daily (1 hour at 37° C) for 4 more days. The candidate flies were balanced and screened by PCR using paired element-specific and genome-specific primers: PB08014: GATCATTAAAACGTGGCCAACTAC (5’ *eag* flanking region e03618) PB08013: CCTCGATATACAGACCGATAAAAC (element specific primer) PB08015: GTGGGGGTTCTTATTCTTCAGTT (3’ eag flanking region f02960).

### Electrophysiology

*Solutions: Nominally Ca^2+^-free HL3.1* (Feng et al. 2004) (in mM): 70 NaCl, 5 KCl, 4 MgCl_2_, 10 NaHCO_3_, 5 Trehalose, 115 sucrose, 5 HEPES; pH adjusted to 7.1-7.2 with NaOH. *0 Ca^2+^ Modified A solution* (in mM): NaCl, 118; KCl, 2; MgCl_2_, 4; Trehalose, 5; Sucrose, 45.5; HEPES, 5; EGTA, 0.5; pH adjusted to pH 7.3-7.4 with NaOH. *Nominally Ca^2+^-free Modified A solution* (in mM): NaCl, 118; KCl, 2; MgCl_2_, 4; Trehalose, 5; Sucrose, 45.5; HEPES, 5; pH adjusted to pH 7.3-7.4 with NaOH. *HEK cell low Ca^2+^ internal patch solution* (in mM): NaCl, 2; K-Gluconate, 130; CaCl_2_, 0.1; MgCl_2_, 2; EGTA, 1; HEPES, 10; pH adjusted to 7.3-7.4 with KOH. *Motor neuron low Ca^2+^ internal patch solution* (in mM): 2 NaCl, 130 K-Gluconate, 0.1 CaCl_2_, 1 EGTA, 0.5 Na-GTP, 4 Mg-ATP, 10 HEPES) adjusted to 285 mOsm with glucose, and pH 7.3-7.4 with KOH.

HEK cell recordings by whole-cell patch were made 48 to 60 hours after transfection. Cells were recorded at 21-22° C in 0 Ca^2+^ modified A solution and a low Ca^2+^ internal patch solution (see solutions above). The appropriate amount of Ca^2+^ was added to the internal solution to yield the approximate calculated free Ca^2+^ concentrations shown in the figure. The Maxchelator program (Chris Patton, Stanford University) was used to calculate the free Ca^2+^ concentrations. The HEK cells were voltage clamped with a Multiclamp 700A amplifier (Axon Instruments, Molecular Devices, Sunnyvale, CA) and currents were filtered at 2.6 kHz and recorded using pClamp 8 software (Axon Instruments, Molecular Devices). Software sampling at 10 kHz. Leak currents were calculated offline by fitting the steady state current vs. voltage relationship from −80 mV to −60 mV. The slope and intercept of the linear fit was used to calculate the leak current at each membrane potential and subtracted from the recorded sustained currents. EAG transfected cells were rejected if the initial resting potential was < 20 mV. The recording for the different EAG isoforms (Figure 1E) were performed as above except that the internal pipette solution was changed to increase EGTA to 5 mM and 4 mM Mg-ATP was added.

**Figure 1.**
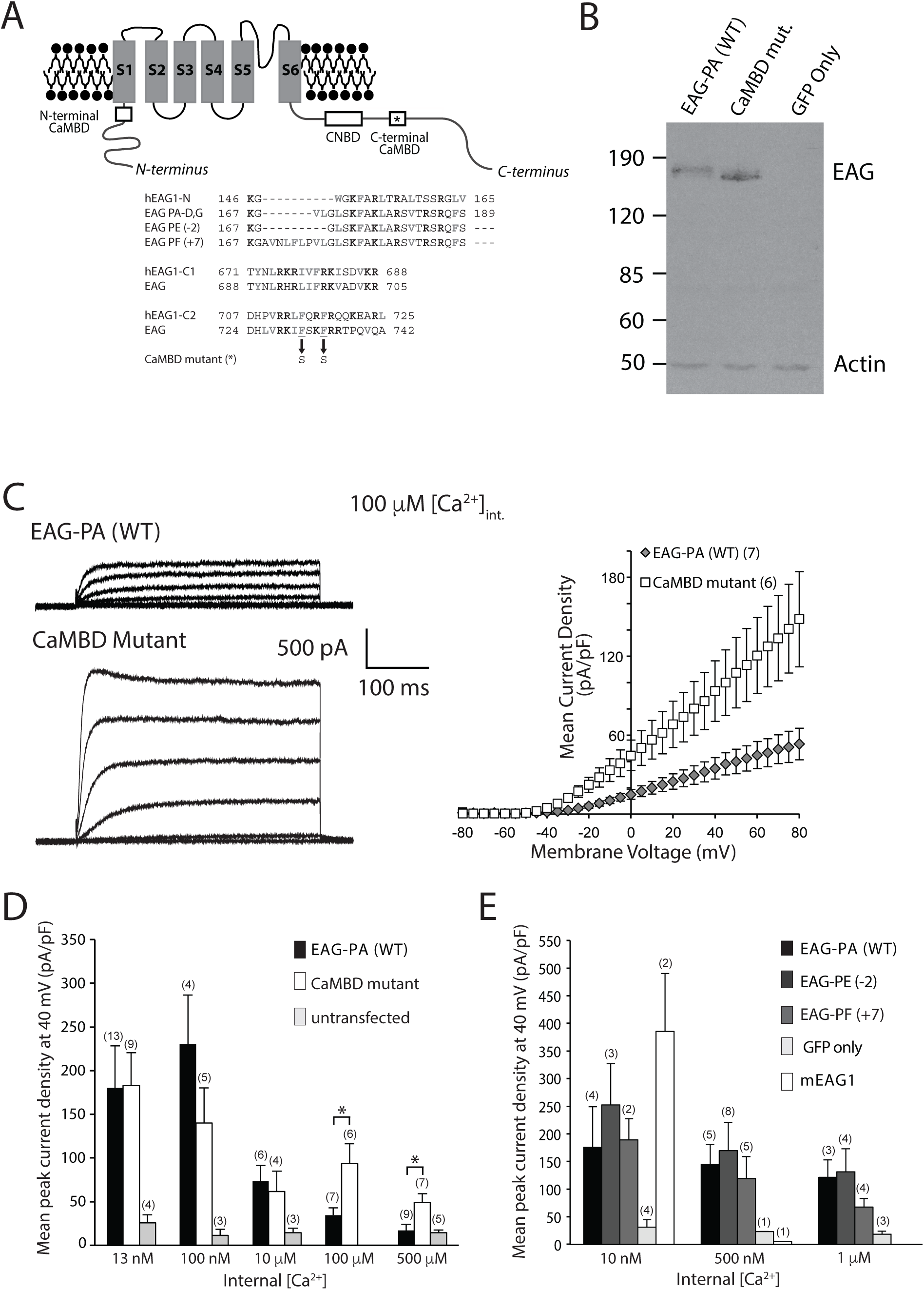
*Drosophila* EAG channels are regulated by Ca^2+^. (A) Cartoon of the fly EAG subunit showing the position of the highly conserved CaM binding domains (CaMBDs, not drawn to scale). Protein sequence alignments of the three fly CaMBDs with human EAG as well as comparison of the different *Drosophila* splice variants in the N-terminal CaMBD. Basic residues are bold black, hydrophobic are bold gray. Arrows indicate the two phenylalanines mutated to serines to disrupt CaM binding in the C2 domain. (B) Western blot of HEK293 cell extracts expressing wild type (EAG-PA) or CaMBD (F232S,F235S) mutant plasmid DNA in combination with an eGFP plasmid. (C) Sample traces of wild type or F232S,F235S mutant EAG currents and I-V relation of EAG current densities recorded under whole-cell patch clamp from HEK293 cells with 100 μM internal Ca^2+^ in pipette solution. (D) Plot comparing mean wild type and CaMBD mutant EAG current densities at various internal Ca^2+^ concentrations at a membrane voltage of 40 mV. Wild type (PA) and CaMBD mutant mean current densities are significantly different at 100 and 500 μM internal Ca^2+^, p = 0.04 for each concentration, Wilcoxon rank sum test. (E) Plot of mean wild type EAG current densities for the N-terminal CaMBD splice variants at 40 mV with mouse EAG for comparison. n represents number of cells recorded. All values are mean ± SEM. There were no significant differences, p = 0.4, 2-factor ANOVA between *Drosophila* splice isoforms across Ca^2+^ concentrations.

Potentials from the neuromuscular junction (NMJ) were recorded in current clamp mode from muscle 6 of wandering third instar (gender specified in fly cross methods) larvae with intracellular electrodes filled with 3 M KCl (20-30 MΩ) in 0.4 mM or 0.2 mM Ca^2+^ HL3.1. Signals were amplified with an Axoclamp 2B amplifier (Molecular Devices, Sunnyvale, CA), digitized with a Digidata 1322A (Molecular Devices), and recorded at 10 kHz with pClamp8 software (Molecular Devices). Muscles with resting potentials more depolarized than −45 mV were not used. The mEJP and nerve evoked recordings were done in 0.2 mM Ca^2+^ HL3.1 except with 2 mM KCl to increase the K+ driving force. For the nerve stimulation experiments, current was injected to keep the muscles at −60 mV to keep a consistent membrane potential for all cells. The motor nerves were stimulated by sucking up the nerve with a glass pipette ~10 μm in diameter at 2-3 times the threshold voltage to first trigger a postsynaptic response. The stimulation was delivered by a model 2100 isolated pulse stimulator (A-M Systems, Sequim, WA) that was triggered by the pClamp software at each recording sweep.

Synaptic currents at the NMJ were recorded from larval muscle 6 of wandering 3^rd^ instar larvae in two-electrode voltage clamp (TEVC) mode with an Axoclamp 2B amplifier and digitized as above. We used 0.2 mM Ca^2+^ HL3.1 solution except that the KCl was adjusted to 2 mM. The 3 M KCl filled voltage sensing electrodes had resistances of 15-18 MΩ, and the current passing electrodes filled with 3 M KCl had resistances of 10-15 MΩ. Muscles were held at −70 mV. Cells were rejected if, after impaling the muscle with both electrodes, the average of the 2 resting potentials recorded at each electrode was more depolarized than −30 mV. Motor nerve stimulation was performed as described in the previous paragraph.

Type Is motor neuron voltages were recorded by whole-cell patch clamp in current clamp mode with a Multiclamp 700A amplifier (Molecular Devices, Sunnyvale, CA). Filtering and sampling as for HEK cells. Larvae were filleted in Ca^2+^-free modified A solution and the brain left intact. Dorsal motor neurons directly below the sheath of the ventral nerve cord were visualized with a 40x water immersion objective and exposed by applying 0.2 % w/v protease (type XIV, MilliporeSigma, St. Louis, MO) in Ca^2+^-free modified A solution with a patch pipette directly around the cluster of neurons. The patch pipettes for recording had resistances of 2-3 MΩ. The identity of the type of motor neuron was based on position in the motor neuron cluster (Choi et al. 2004) and confirmed by the characteristic delay to spike phenotype unique to MNISN-Is motor neurons. Recordings were made first in nominally Ca^2+^-free modified A solution and then the solution was changed to a 1.8 mM Ca^2+^ modified A solution. Current was injected to keep the motor neurons at −60 mV. Cells were rejected if the initial resting potential was more depolarized than −45 mV.

All physiological preparations were superfused with the indicated saline solutions by gravity feed and vacuum suction at 1-2 ml/min. Electrophysiology trace plotting and analysis of membrane current and voltage signals were performed in MATLAB (Mathworks, Natick, MA) using scripts written by PB. Table 1. shows cell properties recorded for each of the data sets in this study.

### Protein analysis

For immunoblots, HEK cells were washed 2x with 1x PBS and then cells were scraped in 500 μl 2x Laemmli sample buffer. The cell suspension was passed through a syringe needle 2-3x to break up the membranes. The homogenate was separated by 7.5% SDS-PAGE, and immunoblotted with an anti-Eag rabbit polyclonal antibody raised against amino acids 1032-1174 of *Drosophila* Eag.

For CaM-agarose purification of WT and CaMBD mutant EAG (which still has an N-terminal CaMBD), 1 ml of flies of each genotype were collected in a 15 ml conical centrifuge tube and frozen on dry ice. The frozen flies were vortexed and dumped onto a sieve to collect the heads that pass through on a fine mesh cooled on dry ice. The heads were put into 1.5 ml eppendorph tubes on dry ice, and then transferred to a Dounce homogenizer (2 ml) on ice. 100 ml of homogenization solution (20 mM HEPES-NaOH pH 7.4, 1 mM EGTA) was added and the heads were homogenized. The homogenate was transferred to a fresh 1.5 ml eppendorph tube and 100 ml additional homogenization solution was added to the homogenizer to wash and collect the remaining homogenate. 200 ml of 2x solubilization buffer (20 mM HEPES-NaOH pH 7.4, 200 mM NaCl, 2% Triton X-100, 1 mM EGTA) was added to the homogenate. Both buffers contained a broad spectrum protease inhibitor, Complete Mini (Roche Diagnostics, Germany).

The tubes of homogenate were rotated at 4° C for 1 hour and centrifuged for 15 minutes (4° C, 14,000 rpm). 3 μl of supernatant was removed from each sample to determine the total starting protein concentration by Bradford protein assay (BioRad).The supernatant was added to 100 μl of sepharose beads conjugated to Calmodulin (equilibrated in same buffers used in solubilization step) along with Ca^2+^ to a final concentration of 2 mM and rotated 1 hour at 4° C. 100 μl of the supernatant (flow through) was removed from each sample before washing the beads in Ca^2+^ containing buffer 3x. 100 μl of 2x Laemmli sample buffer was added to the beads and then the samples were boiled for 10 min. Volumes of samples loaded were based on the protein concentration determined in the Bradford assay. Samples were separated by 7.5% SDS-PAGE, and immunoblotted with an anti-Eag rabbit polyclonal antibody (1:500) raised against amino acids 1032-1174 of *Drosophila* Eag, and Actin antibody, 1:5000 (EMD Millipore).

### Immunohistochemistry

#### Larval brain

Wandering 3^rd^ instar larvae were filleted and pinned out in Sylgard coated 35 x 15 mm petri dishes (3 larvae/dish) containing ice-cold Ca^2+^-free HL3.1 solution. The HL3.1 solution was removed and replaced by fresh ice-cold 4% paraformaldehyde in PBS and incubated at 4° C on an orbital shaker for 4 hours. Fixed preparations were washed 3 × 5 minutes with PBS-TX (0.2% Triton-X-100 in PBS) and then blocked overnight at 4° C in PBS-TX-BSA (0.2% Triton-X-100 and 5% bovine serum albumin in PBS). The head and first few abdominal segments were cut from the rest of the larval body (with the brain attached) and placed into wells of a 60 well minitray (Nunc, Denmark) (1 larva/well) containing rabbit EAG antibody (1:500) in PBS-TX-BSA and mouse Actin antibody, 1:1500 (EMD Millipore) in PBS-TX-BSA. The tissue was incubated at room temperature for 2 hours on an orbital shaker and washed 3 × 5 minutes with PBS-TX. 1:200 Alexa Fluor 488 goat anti-rabbit (Invitrogen, Molecular Probes) and 1:200 Alexa Fluor 680 goat anti-mouse (Invitrogen, Molecular Probes) in PBS-TX-BSA were added to the well and incubated at room temperature for 1.5 to 2 hours. The brain preparations were then washed 3 × 5 minutes and mounted on slides in Vectashield (Vector Laboratories, Burlingame, CA) still attached to the head to keep the brain oriented. Sequential images of the green and red emission signals were taken using the 63x objective on a Leica TCS SP2 confocal microscope. Ten optical sections (1.2 μm each) were taken for each brain using identical settings making sure none of the signals were saturated. The pixel intensities were analyzed using MATLAB (Mathworks, Natick, MA) by averaging the pixel intensity in the same region of the neuropil for each optical section and subtracting the average pixel intensity in the background. The ratio of Green:Red pixel intensities was calculated for each brain.

#### Larval NMJ

Wandering 3^rd^ instar larvae were filleted and pinned out in Sylgard coated 35 x 15 mm petri dishes containing ice-cold Ca^2+^-free HL3.1 solution. The motor nerves were cut carefully and the brain was removed before replacing the solution with ice cold 4% paraformaldehyde in PBS and incubating at room temperature on an orbital shaker for 20 minutes. After a quick rinse in PBS, the prep was washed in PBS for 20 minutes at room temperature on an orbital shaker. Then the larval preparations were washed 3 × 10 minutes at room temperature in PBT (PBS + 0.05% Triton X-100). The larval tissue was then blocked for 1 hour at room temperature in blocking solution (5% heat inactivated goat serum + PBT). Mouse 4F3 anti-DLG antibody (Developmental Studies Hybridoma Bank) was diluted 1:50 in blocking solution and added to the larval preparations and they were incubated overnight at 4° C on an orbital shaker. After 3 x 10 minute washes in PBT, the larval preparations were incubated with goat anti-mouse Alexa-Fluor 488 (Invitrogen) diluted 1:200 in PBS and Alexa-Fluor 594-conjugated goat anti-HRP (1:200, Jackson ImmunoResearch Laboratories) for 2-3 hours at room temperature. Final washes in PBT (5 x 5 minutes). The larval preparations were then mounted on slides in Vectashield (Vector Laboratories, Burlingame, CA). Images of the NMJs were taken sequentially at the appropriate emission spectra on a Leica TCS SP5 confocal microscope. Only boutons associated with DLG staining were counted from muscles 6/7 in segments A2 or A3.

### Cell culture and plasmids

HEK cells were grown in DMEM (Gibco/Life Technologies) supplemented with heat inactivated bovine serum (Invitrogen). Cells were transfected with EAG plasmids (4 μg/100 mm plate) using the Ca^2+^-phosphate method, and co-transfected with GFP (1 μg/100 mm plate). After 6 hours, transfected cells were resuspended and plated onto 35 mm plates coated with poly-L-lysine. 24 hours after transfection, cells were transferred to a 26° C incubator. This was necessary for the expression of functional *Drosophila* Eag channels in HEK cells (Li et al. 2011). HEK cells were transfected with a pcDNA3 vector backbone containing wild type or mutant *eag* cDNA (PA). The C2 CaM binding domain mutations, F232S,F235S were generated by site-directed mutagenesis (Stratagene) using primers: CGTCCGGAAGATATCCTCCAAATCTCGTCGCACTCCGC (forward) and GCGGAGTGCGACGAGATTTGGAGGATATCTTCCGGACG (reverse). The correct sequence of the plasmids was verified over the entire length of the cDNA. The splice isoforms EAG-PE and EAG-PF were constructed from the EAG-PA plasmid described above with the Gibson cloning method using 5 primers:

F10: GACTCACTATAGGGAGACCCAAGCTTGGTACCGAGCTCGGATCC

R11: GCCAATTTGGCGAATTTCGAGAGACCTCCCTTCGTGTCCTCGCTGTCG

R14:GCGAATTTCGAGAGACCAGGGAGAAACAGGTTGACGGCAAAACTCCCTTCGTGTCCTCG

F12: GTCTCTCGAAATTCGCCAAATTGGCAAGATCAGTGACACG

R13: TTCTCGTTGTCTGTCTCCGCGGCGACATTGCCAAAGCCCACC 3 PCR reactions were performed. PCR a-F10:R11, PCR b-F10:R14, PCR c-F12:R13. Region of editing was checked by sequencing.

### Calcium Imaging of Adult *Drosophila* Neurons

Homozygous *eag* CaMBD17 mutant and *eag* WT12 control female flies were crossed with transgenic flies carrying *GH146-GAL4* (Bloomington Stock # 30026) and *UAS-GCaMP6f* (Bloomington Stock # 52869). Male progeny were screened for transgene expression by eye color and housed at 25°C for 5-10 days post eclosion. Experimental animals were anesthetized with ice, and brains were dissected in Adult Hemolymph-Like (AHL) saline (Wang et al. 2003). *Ex vivo* brains were secured to a Sylgard-lined perfusion chamber filled with AHL using bent tungsten pins.

The antennal lobe (AL) of the brain was located by morphology using an Olympus BX51WI microscope and 40X/0.8NA objective. A FHC concentric bipolar electrode (CBAEB50) was positioned in the center of the AL using a manual micromanipulator. During each experimental trial, 4 electrical stimulus trains were delivered to the AL by an A.M.P.I. Master-8 pulse generator with a 20 second inter-train interval. The stimulus trains were 20 pulses with a 1ms pulse duration at a 100Hz pulse rate. The stimulus voltage was raised on each trial; 3, 5, 7, and 9 volt trials were performed on each brain.

The GCaMP6f response of the *GH146* terminals in the LH was measured using a Hammamatsu Orca-ER camera controlled by Micro-Manager. Images were acquired at a 300 msec frame interval. Stimulus delivery and image acquisition were synchronized using a National Instruments USB-6212 DAQ and a custom MATLAB program. Brains were continuously perfused with room temperature (20-22°C) AHL during the experiment. All brains responded to the high voltage (9V) stimulus, no brains were excluded, and the stimulus electrode was not repositioned after the experiment began.

Imaging data were analyzed using a custom MATLAB program. The program aligned the image time series to correct drift using the Image Processing Toolbox. An analysis region of interest (ROI) and background (BG) region were manually defined, the per-frame mean fluorescence of the ROI and BG were calculated, and the BG corrected fluorescent time series was calculated. The ΔF/F was calculated for each frame *i* using the formula: ΔF_*i*_ / F = (F_*i*_ - F_1-20_) / F_1-20_. The peak change in ΔF/F was calculated as the ΔF/F of the stimulus frame minus the ΔF/F of the frame before the stimulus. The sustained level of ΔF/F was calculated as the mean ΔF/F of frames 1-5 seconds after the stimulus minus the ΔF/F of the frame before the stimulus. Frame time-stamps were used to generate a mean peri-stimulus ΔF/F of each trial. The timing of each frame varied by several milliseconds due to software delays so, for display, the data was re-sampled to a uniform 10 Hz time base using shape-preserving piecewise cubic interpolation.

### Modeling

#### Characterization of EAG channels

Time series data of currents in HEK cells expressing EAG channels during voltage clamp were used to characterize the activation functions and kinetics of EAG channels. Peak currents in HEK cells as a function of voltage were subtracted from peak currents in EAG-expressing HEK cells to isolate the currents due to EAG channels alone. Leak currents were then subtracted from this I-V curve, which was then converted to a curve of conductance *vs.* voltage using a reversal potential *E*_*K*_ = −106.17mV. EAG channels were modelled using the Hodgkin-Huxley framework (Hodgkin and Huxley 1952). The current due to EAG channels is given by

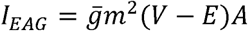

where 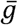 is the specific conductance of the channels, *m* is the activation variable, *E* is the reversal potential and *A* is the surface area of the compartment. *m* is a time varying variable determined by

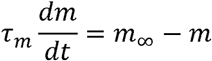

where *m*_∞_ and *τ_m_* are functions of the membrane potential *V* that are determined by fitting functions to plots of conductance *vs*. voltage and time-series of conductance at various holding voltages. Since the activity of EAG channels also depends on the intracellular Ca^2+^ concentration, the process was repeated for multiple Ca^2+^ concentrations to obtain functions that depended on both the membrane potential and the Ca^2+^ concentration. For these channels, we used

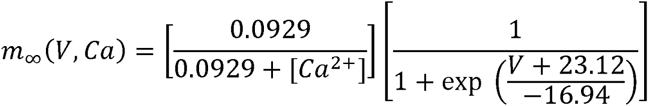

for wild-type EAG channels and

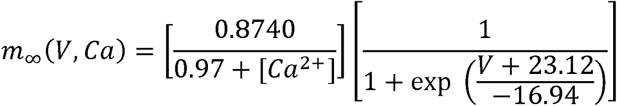

for CaMBD mutant EAG channels. Note that the mutant and wild-type channels differ in their Ca^2+^ sensitivity, but not in their voltage sensitivity (Figure 1C). For both wild-type and mutant EAG channels, we found that the timescale of activation was not a strong function of Ca^2+^ concentration; so we used

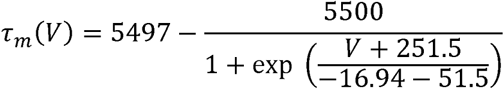

In these functions, *V* is in units of mV, *τ*_m_ is in ms, and [*Ca*^2+^] is in *μ*M.

*Model of presynaptic terminal*. Because EAG is inhibited only at relatively high levels of intracellular Ca^2+^, we reasoned that a possible zone where EAG could be inhibited is the microdomain close to voltage-gated Ca^2+^ channels (VGCCs) at the presynaptic terminal. We modelled the presynaptic terminal using a three-compartment model: one for the invading action potential, another for the Ca^2+^ microdomain, and a third for parts of the pre-synaptic terminal far from VGCCs. The Ca^2+^ microdomain had voltage-gated Na^+^ channels, K^+^ channels, EAG channels, and Ca^2+^ channels. The other compartment had the same channel profile, except for Ca^2+^ channels. All compartments were electrically coupled to each other with a conductance of 0.1 nS.

The activation functions and kinetics of Na^+^ channels were determined using fits to data from embryonic *Drosophila* neurons (O’Dowd and Aldrich 1988); K^+^ channels from *Drosophila* photoreceptors (Hardie 1991) and Ca^2+^ channels from *Drosophila* embryonic muscles (Hara et al. 2015).

The model was integrated using the exponential Euler method (Dayan and Abbott 2001) using a general-purpose neuron simulator written in C++, that we have made freely available at https://github.com/sg-s/xolotl/.

#### Characterization of effect of EAG on action potential width

To study the effect of WT and mutant EAG on the action potential waveform, we set up a single compartment model with Na^+^, K^+^, Ca^2+^ and EAG channels as described in the previous section. The Ca^2+^ dependence of the EAG channel was constrained by data from recordings of WT of mutant EAG channels. In Panel A, the “EAG null” trace was generated without any EAG channels in the single compartment model. In panels B-D, all parameters except the one being varied on the X-axis were kept constant at the reference value (g_Na_ = 500 μS/mm^2^, g_K_ = 60 μS/mm^2^, g_EAG_ = 300 μS/mm^2^, g_Ca_ = 20 μS/mm^2^, g_Leak_ = 1 μS/mm^2^).

#### Code availability

A script to reproduce Figure 2 and 6, a toolbox to interactively vary parameters in the model, and scripts to estimate gating functions of EAG from raw data are available at https://github.com/marderlab/eag.

**Figure 2.**
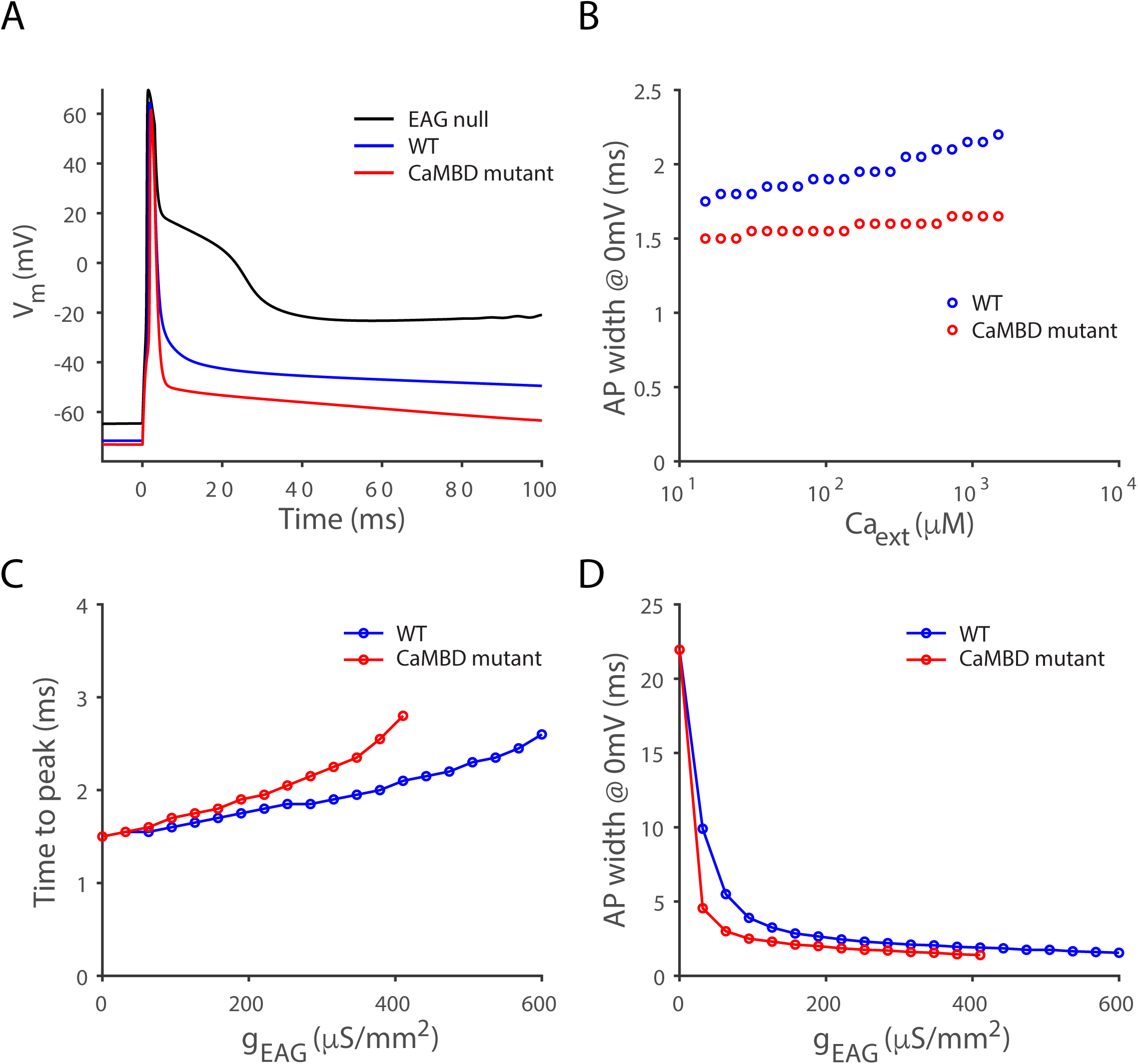
Comparison of the effects of *eag* null and CaMBD mutations on action potential properties in a one-compartment model of *Drosophila* presynaptic terminal. (A) Evoked potentials in response to a 3 msec depolarization for WT EAG (blue), *eag* null mutant (black) and CaMBD mutant (red). (B) Effect of genotype on action potential (AP) width at 0 mV. The presence of EAG allows Ca^2+^ to influence AP width. (C) Time to peak current is affected by both the amount of EAG and its Ca^2+^ sensitivity. (D) AP width is affected by EAG Ca^2+^ sensitivity over a limited range of current densities.

### Learning and memory assays

Appetitive conditioning protocol was as described previously (Liu et al., 2012). A group of ~50 flies in a training tube alternatively received 10% 3-octanol (OCT; Merck) and 10% 4-methylcyclohexanol (MCH; Sigma-Aldrich) for 1 min for immediate memory or for 2 min for 24 h memory. Flies were loaded into CS+ tubes containing 2M dried sucrose filter papers with one odor presentation and then CS-tubes without sucrose papers with the other odor. After the retention time being tested (2 min or 24 h), the trained flies were allowed to choose between OCT and MCH for 2 min in the T-maze. A learning index was then calculated by taking the mean performance indices (PIs) of the two reciprocally trained groups. Each PI was calculated by the differences of flies choosing CS+ and CS-divided by the total number of flies. To control whether the learning deficits were due to lack of ability to sense sucrose or odors, flies were allowed to choose between sucrose+ and sucrose-tubes for 2 min or between one odor (OCT or MCH) and air for 2 min. Sucrose preference index or odor avoidance index was calculated, respectively.

### Statistics

#### Electrophysiology

Wilcoxon Rank Sum, unpaired Student’s t-test and ANOVA done in MATLAB (MathWorks, Natick, MA) or Excel (Microsoft).

#### Memory

statistical analyses were performed using GraphPad Prism 7. The Wilks-Shapiro test was used to determine data normality. Normally distributed data were analyzed with unpaired t-test, and data which did not pass the normality test were analyzed with a Mann-Whitney test. Data are presented as mean ± SEM. Difference between groups were considered significant at p < 0.05. * = p < 0.05; ** = p < 0.001; *** = p < 0.0001.

## Results

### *Drosophila* EAG is inhibited by high [Ca^2+^]_i_

Human EAG has three calmodulin (CaM) binding sites: one N-terminal and two C-terminal (C1 and C2) (Ziechner et al. 2006) and is inhibited by nM levels of Ca^2+^ (Schonherr et al. 2000). Previous work on *Drosophila* EAG, in which it was compared to hEAG1 in *Xenopus* oocytes, suggested that the fly channel was not regulated by Ca^2+^ (Schonherr et al. 2000). Intrigued by the fact that all of these sites appear to be conserved in *Drosophila* (Figure 1A), we revisited the issue of regulation by Ca^2+^ by expressing wildtype (WT) EAG, or a C-terminal CaMBD mutant form (F732S,F735S, Sun et al. 2004), in HEK293 cells and recording currents with varying amounts of Ca^2+^ in the pipette internal solution. The C-terminal mutation should block all CaM-dependent regulation, since inhibition has been suggested to depend on CaM bridging the N-and C-termini (Whicher and MacKinnon 2016). Mutant and WT channels were expressed at comparable levels (Figure 1B), though there can be variability in molecular weight that is likely due to differences in glycosylation (see Fig 3B and Ramos Gomes et al. 2015). GFP was co-expressed to identify transfected cells.

**Figure 3.**
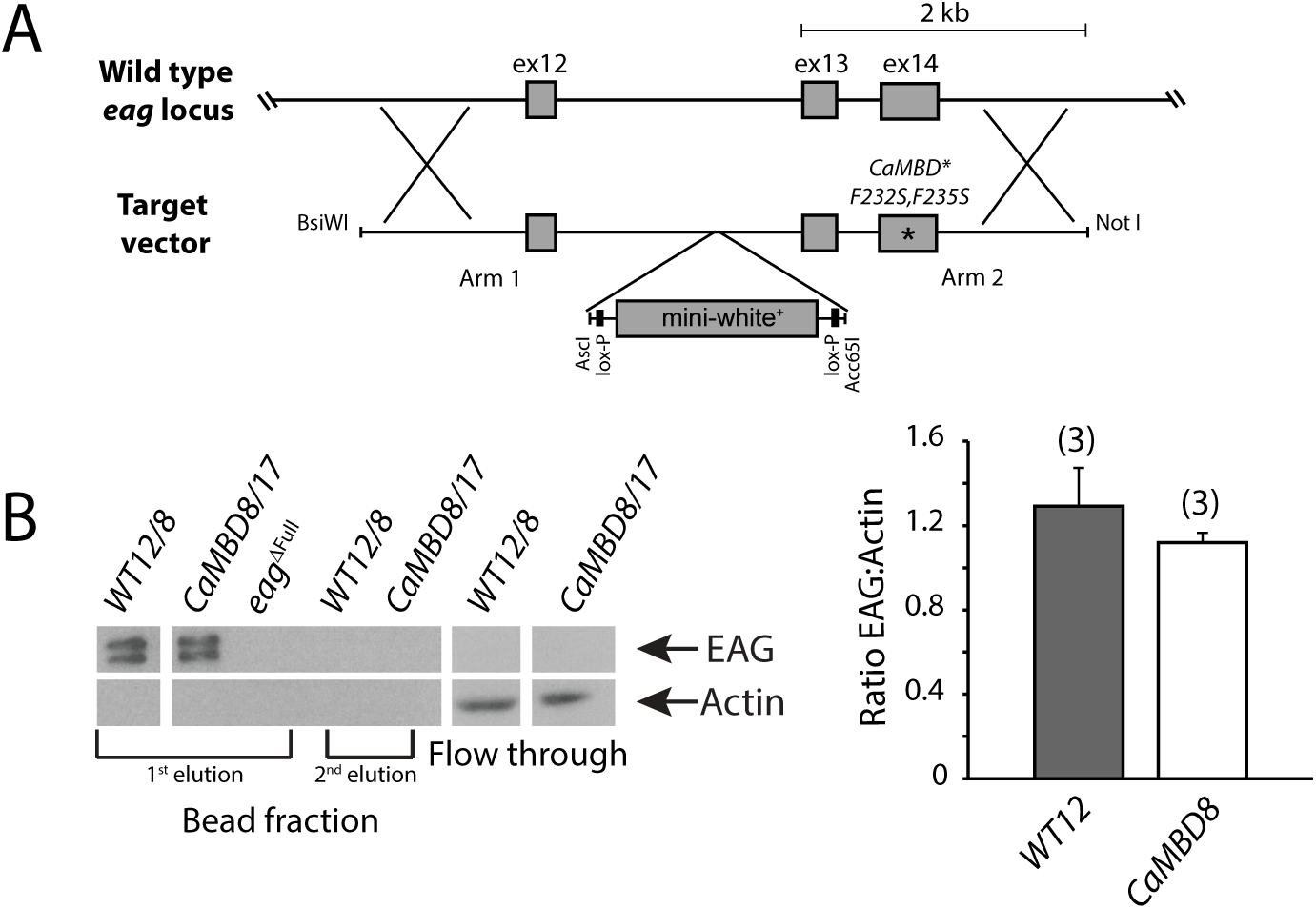
Characterization of F232S,F235S CaMBD homologous recombination mutant *Drosophila*. (A) Schematic of the targeted region (exons 12-14) of the *Drosophila eag* locus and the target vector used to generate the CaMBD mutants with F232S,F235S mutations. The control line used the same target vector without the F232S,F235S mutations. (B) Western blot (left) from a CaM bead pull-down of fly EAG from whole head extract of CaMBD mutant, WT control and the *eag*^*ΔFull*^ null allele. The flow through and 2^nd^ bead elutions were run to confirm all of EAG was bound and released in the 1^st^ elution. Flow through shows actin levels, indicating loading was comparable for these samples. Larval brain immunohistochemistry (right) with EAG antibody. Plot of the mean pixel intensity of the neuropil normalized to Actin immunostaining for control and CaMBD mutant larvae. There is no significant difference in the normalized pixel intensities for WT and CaMBD mutants, p = 0.4, unpaired Student’s t-test. n represents number of brains. Values are mean ± SEM.

As previously reported (Schonherr et al. 2000), *Drosophila* EAG was not inhibited at or below 1 μM [Ca^2+^]_i_. Only when [Ca^2+^]_i_ was increased to high μM levels, did we see significant inhibition. Figure 1C (left panel) shows currents recorded in the presence of 100 μM internal Ca^2+^. Currents from cells transfected with WT EAG are clearly smaller compared to those from cells expressing EAG carrying mutations blocking CaM-binding to the major C-terminal site, reflecting inhibition by Ca^2+^. The voltage-dependence of the channel is unaffected by this mutation (Figure 1C, right panel). A dose-response relationship for [Ca^2+^]_i_ is shown in Figure 1D. In spite of high variability, it is clear that WT EAG current density approaches that of untransfected cells at high [Ca^2+^]_i_ while the CaMBD mutant has substantial residual current even at almost mM levels of Ca^2+^ in the pipette.

It is interesting to note that although the C-terminal CaMBD should completely block the gating mechanism proposed by Whicher and Mackinnon (2016), the CaMBD mutant does show substantial inhibition at high Ca^2+^ levels. Whether this inhibition is due to some action of CaM mediated by the N-terminal CaMBD alone or whether it is due to some other Ca^2+^-dependent regulatory process (e.g. activation of a cyclase or phosphorylation by CaMKII Wang et al. 2002) is unknown.

### N-terminal splicing does not substantially alter the Ca^2+^ sensitivity of EAG

Examination of genomic data for the *eag* gene revealed that the N-terminal CaMBD, which is a critical partner of the C-terminal site for Ca^2+^-dependent gating (Whicher and MacKinnon 2016), is located at an exonic junction that is alternatively spliced (Attrill et al. 2016, Figure 1A). Since CaM is known to bind amphipathic helices, variation at this region is potentially important for the affinity of CaM-binding and might explain the published Ca^2+^-insensitivity of the canonical “WT” EAG, which corresponds to the PA splice variant (Drysdale et al. 1991) curated on FlyBase, if PA produced a dysfunctional N-terminal binding domain. The splice site found in PA is also present in the PD and PG proteins, but the PE splice form has a 2 amino acid deletion, making it more like the mammalian EAG1 protein, while the PF variant has an additional 7 amino acids inserted into the predicted amphipathic helix. To determine if splice variation at this site substantially alters the Ca^2+^-sensitivity of EAG, we transfected each of the splice forms into HEK293 cells and assayed their response to changes in [Ca^2+^] (Figure 1E). Neither PE nor PF were more sensitive to Ca^2+^ than the original PA isoform and all were significantly less sensitive than mEAG1, which was completely inhibited at 500 nM [Ca^2+^]_i_. We conclude that either the N-terminal splice variants all display the same low affinity for Ca^2+^/CaM or the functional sensitivity of *Drosophila* EAG to inhibition by Ca^2+^ is set by low affinity of the C-terminal CaMBDs since it is unchanged by variation in the N-terminal domain.

### Modeling a role for Ca^2+^-dependent inhibition of EAG

The biological role of Ca^2+^-dependent inhibition of EAG channels is not easily intuited, since K^+^ channels are usually thought to function to suppress excitability. Indeed all previous work on *Drosophila eag*, using mutants that eliminate or reduce EAG currents, are consistent with this idea (starting with Ganetzky and Wu 1983; Wu et al. 1983). EAG channels are thought to contribute to repolarization and their loss broadens the neuronal action potential, prolongs release at the larval neuromuscular junction (NMJ) in flies and induces spontaneous depolarization of the presynaptic terminal (Griffith et al. 1994a).

To aid in thinking about how Ca^2+^-dependent inhibition of EAG might affect the shape of the action potential, we constructed a single-compartment, conductance-based model of the presynaptic terminal. Data from our HEK cell experiments were used to model EAG, and literature values were used for Na^+^, K^+^ and Ca^2+^ channels. We modeled three situations: WT (EAG with Ca^2+^-sensitivity measured in WT), *eag* null (no EAG) and CaMBD mutant (WT levels of EAG, but with Ca^2+^-sensitivity as measured from the CaMBD mutant in HEK cells). Figure 2A shows the effects of a 3 msec depolarization on membrane potential. Compared to the WT case, the *eag* null mutant has a very prolonged and broadened action potential, qualitatively similar to what has been seen *in vivo* in recording from animals with reduced amounts of EAG. In contrast, the CaMBD mutant EAG produces a marked narrowing of the action potential, consistent with a failure of Ca^2+^ influx to inhibit the current. Additional metrics for the action potential waveform suggest that the Ca^2+^-dependent inhibition of EAG may serve to broaden spikes (Figures 2B-D).

The low sensitivity of EAG to Ca^2+^/CaM inhibition is notable. How EAG would respond in the context of the cell is not clear. Bulk cytoplasmic Ca^2+^ is modulated in the nM to low μM range in neurons. The HEK cell results show that at very high levels of intracellular Ca^2+^, the CaMBD mutant EAG channel has a higher conductance than the WT channel (Figure 1D). The only cytoplasmic domains that have levels of Ca^2+^ (mM) that are this high are near the mouth of open Ca^2+^ channels (Parekh 2008). The affinity of EAG for Ca^2+^/CaM suggests that it would only be inhibited if it were localized to this type of Ca^2+^ channel microdomain *in vivo*, meaning that regulation may be more complicated in a real neuron.

### Construction and characterization of a CaMBD mutant at the endogenous *eag* locus

To determine if there were a physiological role for Ca^2+^-dependent inhibition of EAG, we engineered the C-terminal CaMBD mutations characterized above into the endogenous *eag* locus by ends-out homologous recombination (Figure 3A). This mutant is qualitatively different from previously characterized loss-of-function or null *eag* mutants in that it does not reduce the EAG K^+^ current, rather it allows it to persist when Ca^2+^ is high by blocking its inhibition. Multiple alleles containing the F732S,F735S mutations were recovered. WT strains, made in an identical manner with WT recombination arms were also generated. Like the mutant alleles, these WT strains contain a residual 76 bp loxP site located in an intron. These strains were used in all experiments as controls due to their similar genetic background.

Biochemical analysis of adult head protein from CaMBD and WT flies by partial purification on CaM-agarose and immunoblotting shows that EAG protein is present in comparable amounts in both mutant and WT control alleles (Figure 3B, left panel). Quantitative immunohistochemistry of larval brain neuropil (Figure 3B, right panel) confirms this. EAG is normally present at high concentration in neuropil and can be seen in axons and at the NMJ in larvae (see Sun et al. 2004, Fig 1B for immunohistochemistry of native EAG) and the CaMBD mutation did not change this distribution (data not shown).

### Third instar NMJs in CaMBD mutants are morphologically normal and have no spontaneous activity

The NMJ of the third instar larva has been extensively used to assay excitatory synaptic structure and function. *eag* null mutants, with reduced EAG K^+^ conductance, have a characteristic constellation of phenotypes in this preparation which include increases in the number of synaptic boutons (Budnik et al. 1990) and a robust hyperexcitability phenotype in larval motor neurons seen both postsynaptically at isolated NMJs (Drysdale et al. 1991; Ganetzky and Wu 1983; Griffith et al. 1994b) and at the soma (Srinivasan et al. 2012). To determine if CaMBD mutants had morphological abnormalities we stained third instar NMJs with antibodies to HRP and DLG to visualize presynaptic and postsynaptic structures. NMJs from CaMBD and WT strains were indistinguishable (Figure 4A) and there was no significant difference in bouton number.

**Figure 4.**
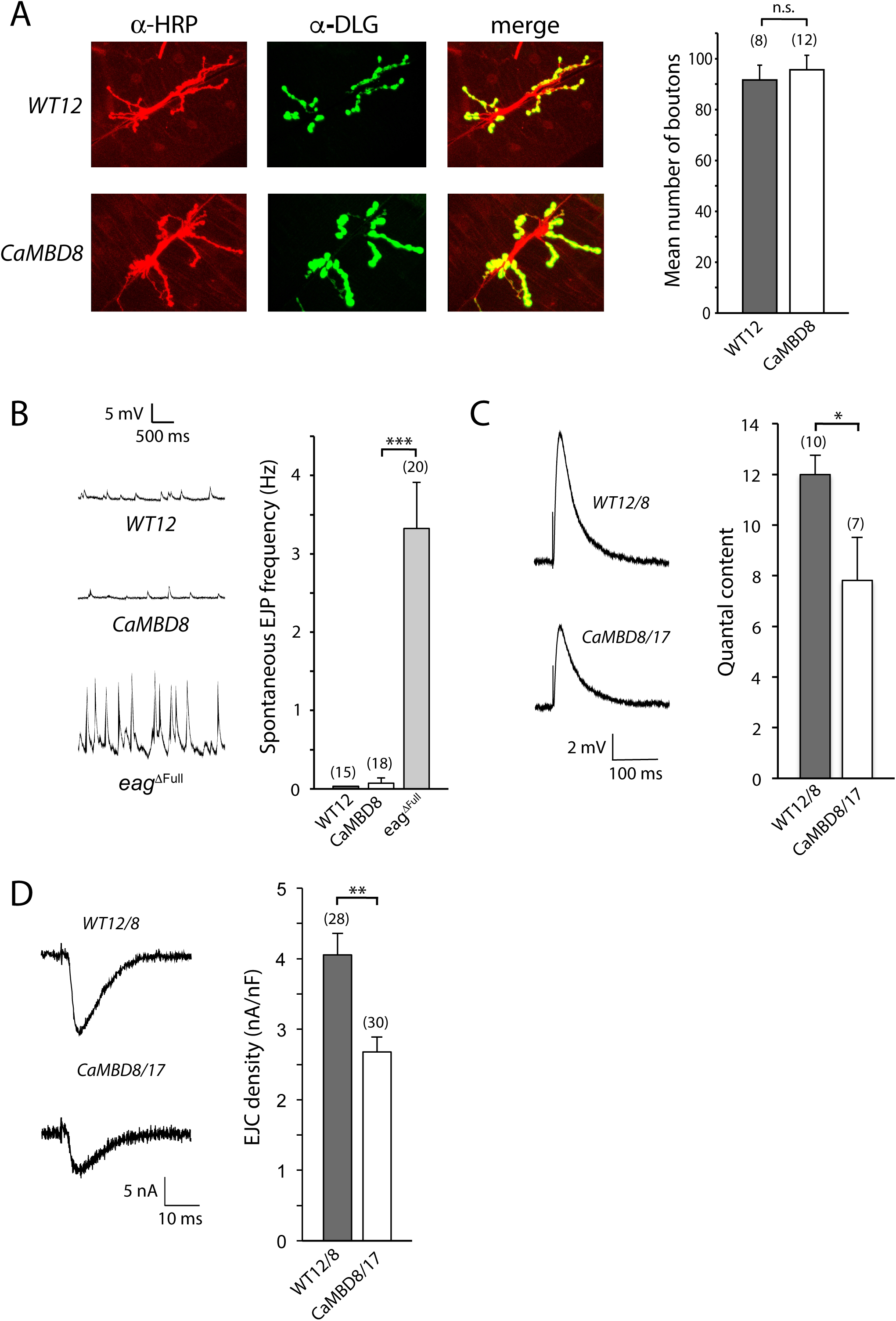
Characterization of the CaMBD mutant larval neuromuscular junction (NMJ). (A) Sample images from immunohistochemistry of WT and CaMBD mutant larval neuromuscular junctions (muscles 6/7) with HRP and DLG antibodies to compare morphology of motor neuron boutons. Plot of number of boutons at the muscle 6/7 NMJs for WT and CaMBD mutants. There is no significant difference in the number of CaMBD mutant boutons compared to WT, p = 0.6, unpaired Student’s t-test. n represents number of NMJs from 4 larvae each for WT control and CaMBD mutant. (B) Sample traces of spontaneous excitatory junctional potentials (EJPs) in WT, CaMBD mutant and a molecular null of *eag* recorded in 0.4 mM external Ca^2+^. Plot of the mean spontaneous EJP frequencies. The CaMBD mutants are significantly different from the *eag* full deletion, p = 1.1 × 10^−7^, Wilcoxon rank sum test. n represents number of cells from 5 WT larvae, 6 CaMBD mutant larvae, and 7 full *eag* deletion larvae. (C) Sample traces of stimulated EJPs and plot of mean quantal content in 0.2 mM external Ca^2+^. The CaMBD mutants had a significantly smaller quantal content, p = 0.03, Wilcoxon rank sum test. n represents number of cells from 9 larvae for WT and 6 larvae for the CaMBD mutants. (D) Sample traces of stimulated excitatory junctional currents (EJCs) and plot of mean EJC amplitudes recorded in 0.2 mM external Ca^2+^ and normalized to muscle size. The mean current density of the CaMBD mutant EJCs were significantly smaller than controls, p = 4.7 × 10^−4^, unpaired Student’s t-test. n represents number of larvae (1 cell recorded per larva). n.s. = not significant. Values are mean ± SEM.

We also performed current clamp recordings from muscle 6/7 to ask if the CaMBD mutant was functionally normal. Baseline recordings from CaMBD and WT control animals showed no significant level of spontaneous excitatory junctional potentials (EJPs, Figure 4B). In contrast, recordings from NMJs of *eag*^*ΔFull*^, a molecular null mutant, had high levels of spontaneous activity. In experiments in which we evoked release by nerve stimulation in normal saline, we did not see any supernumerary EJPs in the CaMBD mutant (P.B., data not shown).

### CaMBD mutant third instar larvae exhibit decreases in the amplitude of evoked EJPs

The EJPs seen in unstimulated *eag* null mutants are the result of spontaneous depolarization of the presynaptic nerve terminal and synchronous Ca^2+^-dependent release of many vesicles. To determine if there were changes in spontaneous Ca^2+^-independent release of single vesicles in the CaMBD mutants, we recorded miniature excitatory junctional potentials (mEJPs). Neither the rate nor the amplitude of this class of events was changed in CaMBD animals compared to WT controls (mean mEJP frequency: WT12/8, 1.92 ± 0.21 Hz; CaMBD8/17, 1.46 ± 0.15 Hz, p = 0.09. Mean mEJP amplitude: WT12/8, 0.85 ± 0.04 mV; CaMBD8/17, 0.83 ± 0.05 mV, p = 0.8). This indicates that the probability of release for individual vesicles and the quantal postsynaptic response to glutamate are both unaffected by loss of regulation of EAG by Ca^2+^.

We also assayed evoked release by stimulating the motor nerve with a suction pipette and recording activity in the muscle. We saw a significant decrease in the amplitude of stimulated EJPs in the CaMBD mutants and a reduced quantal content (Figure 4C). EAG is also expressed in larval muscles (Zhong and Wu 1991; 1993), so to make sure that activation of the EAG channels in the postsynaptic muscle did not confound the amplitude of potentials recorded in the muscle, we voltage clamped the muscle and recorded excitatory junctional currents (EJCs). As seen in Figure 4D, CaMBD mutants had significantly smaller EJC amplitudes than controls. There was no significant difference in the decay of the CaMBD mutant EJCs compared to WT controls (WT: 10.2 + 0.5 ms, CaMBD mutant: 10.8 + 1.8 ms, p = 0.8, unpaired Student’s t-test.

### CaMBD domain mutants have reduced presynaptic Ca^2+^ influx

The CaMBD mutant’s reduction in quantal content and synaptic currents is consistent with a gain of EAG function (i.e. more K^+^ current) when Ca^2+^/CaM is not able to inhibit EAG currents. As seen in Figures 1C and D, the F732S,F735S mutation leads to larger EAG currents when internal Ca^2+^ is at high concentrations. This could cause more effective repolarization of activated presynaptic terminals and an early termination of Ca^2+^ influx, which would be predicted to decrease neurotransmitter release. To directly test the effects of the CaMBD mutation on Ca^2+^ influx, we expressed GCaMP6, a genetically encoded Ca^2+^ indicator (Chen et al. 2013) in olfactory projection neurons. These neurons have well-separated axonal and dendritic processes that allow clear visualization of the presynaptic terminals in the lateral horn. These neurons can be directly stimulated by applying current to the antennal lobes with a bipolar stimulating electrode.

Figure 5 shows GCaMP responses in WT and CaMBD mutant terminals with different levels of stimulation. Ca^2+^ influx increases with stimulation strength, but CaMBD mutant sustained levels are lower than WT at all voltages. Interestingly, the form of the WT Ca^2+^ response changes at high stimulation levels. The peak appears to plateau and a shoulder appears that increases the width of the response. These data are consistent with the effect of the CaMBD mutant having more repolarizing K^+^ current than WT and terminating the Ca^2+^ influx from voltage-gated Ca^2+^ channels more effectively.

**Figure 5.**
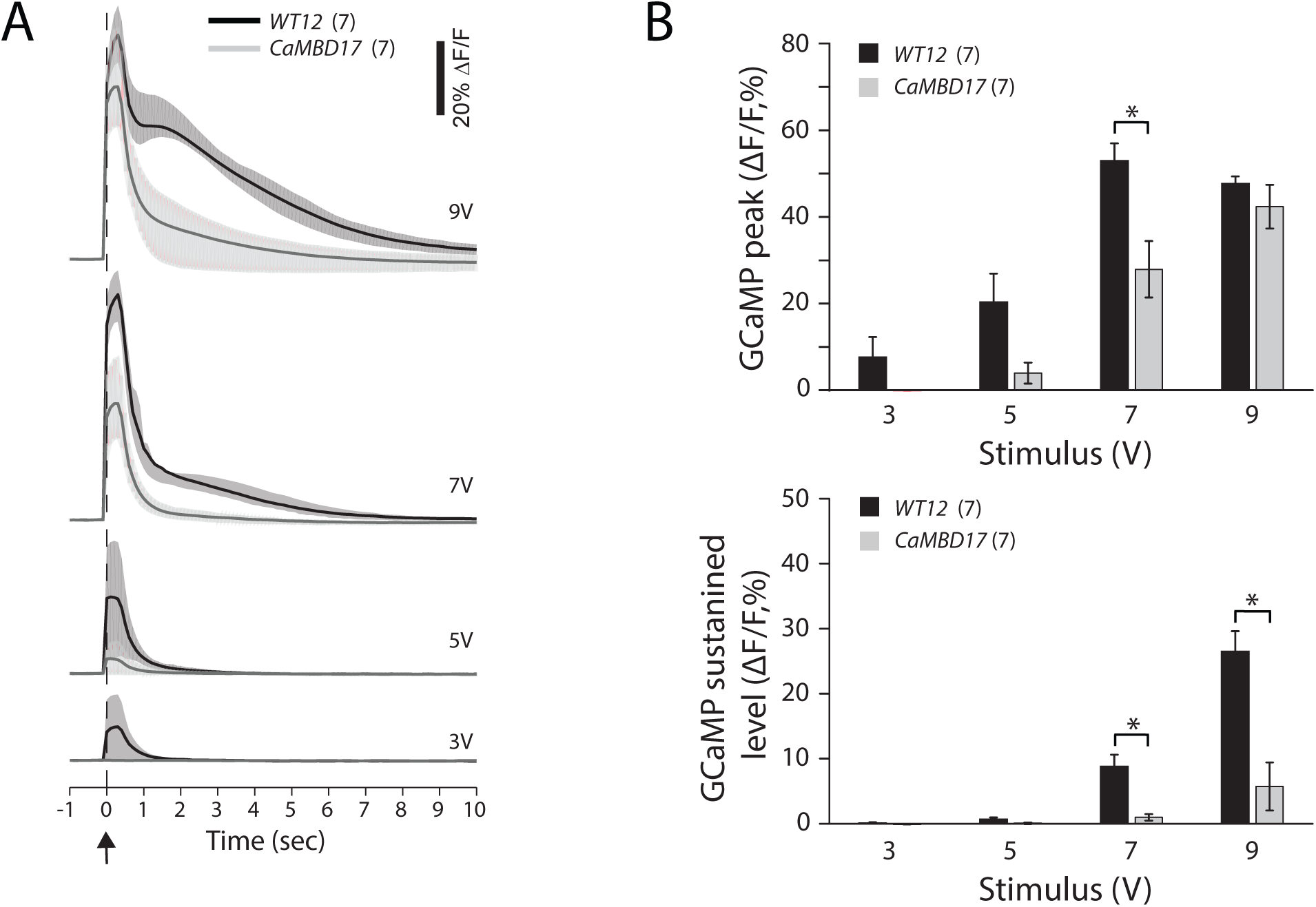
CaMBD mutants have reduced evoked presynaptic Ca^2+^ influx. (A) Mean ΔF/F (%) traces from imaging of GCaMP6 responses in the lateral horn axon terminals of *GH146-GAL4+* neurons from adult brains before, during, and after electrical stimulation of the antennal lobe. Shaded areas represent the 25^th^/75^th^ percentile bounds. Arrow indicates time of stimulus (20 X 1 msec, 100 Hz). (B) Bar graphs of mean peak change in ΔF/F (top) and sustained level of Δ F/F (bottom) for each stimulus amplitude. With a 7 V stimulus, the mean peak change was significantly lower in CaMBD mutants compared to controls, p = 0.03, and the sustained level was significantly lower in CaMBD mutants at 7 V, p = 0.004, and 9V, p = 0.001; unpaired Student’s t-test with Bonferroni correction for multiple comparisons. n represents the number of brains. Values are mean ± SEM.

### A multi-compartment model of the presynaptic terminal recapitulates changes in the width of action potential and duration of the Ca^2+^ influx seen in the CaMBD mutant

The fact that Ca^2+^ influx appears to be blunted in the CaMBD mutant supports the idea that this mutation produces more repolarizing K^+^ current. However since the sensitivity of *Drosophila* EAG to Ca^2+^ is very low, it is possible that this only matters if EAG is present close to areas of high local Ca^2+^. In presynaptic terminals, the voltage-gated Ca^2+^ channels that control neurotransmitter release are highly localized to the active zone, a molecular complex where vesicle fusion occurs (Kawasaki et al. 2004). In contrast, *Drosophila* EAG immunohistochemistry (Gillespie and Hodge 2013; Sun et al. 2004) suggests that EAG is widely distributed, including in areas where it would never see Ca^2+^ levels high enough to cause inhibition. To determine if our electrophysiological and imaging results were consistent with the presence of two distinct pools of EAG, one that could be inhibited by Ca^2+^ and a second that was insensitive even in WT animals, we built a model of the presynaptic terminal that contained pools of EAG that were both inside and outside of Ca^2+^ microdomains (Figure 6). Using parameters obtained from our HEK cell experiments, we simulated the invasion of an action potential into the presynaptic terminal and studied its effects in regions corresponding to the two pools of EAG. Consistent with our results at the NMJ and in the adult brain, we found that action potentials were narrower (Figure 6B), and Ca^2+^ influx smaller (Figure 6D), in CaMBD mutants compared to WT. Since EAG channels outside Ca^2+^ microdomains were inhibited to a smaller degree, membrane repolarization occurred more quickly compared to inside Ca^2+^ microdomains (Figure 6C).

### Loss of Ca^2+^-dependent inhibition of EAG is associated with increased somatic excitability

While the decrease in presynaptic release at the NMJ appears to be due to a decrease in excitability and Ca^2+^ influx, direct recordings from terminals are difficult, so we carried out whole-cell patch clamp of identified MNISN-Is neurons in the ventral ganglion to more closely examine the relationship between membrane potential and firing in the CaMBD mutant. Figure 7A shows representative traces of CaMBD and WT control neurons with increasing current injection. CaMBD neurons fire earlier and at a higher rate than that of WT controls. The complete F/I curves are shown in Figure 7B. These data demonstrate that the CaMBD mutants are somatically hyperexcitable.

**Figure 6.**
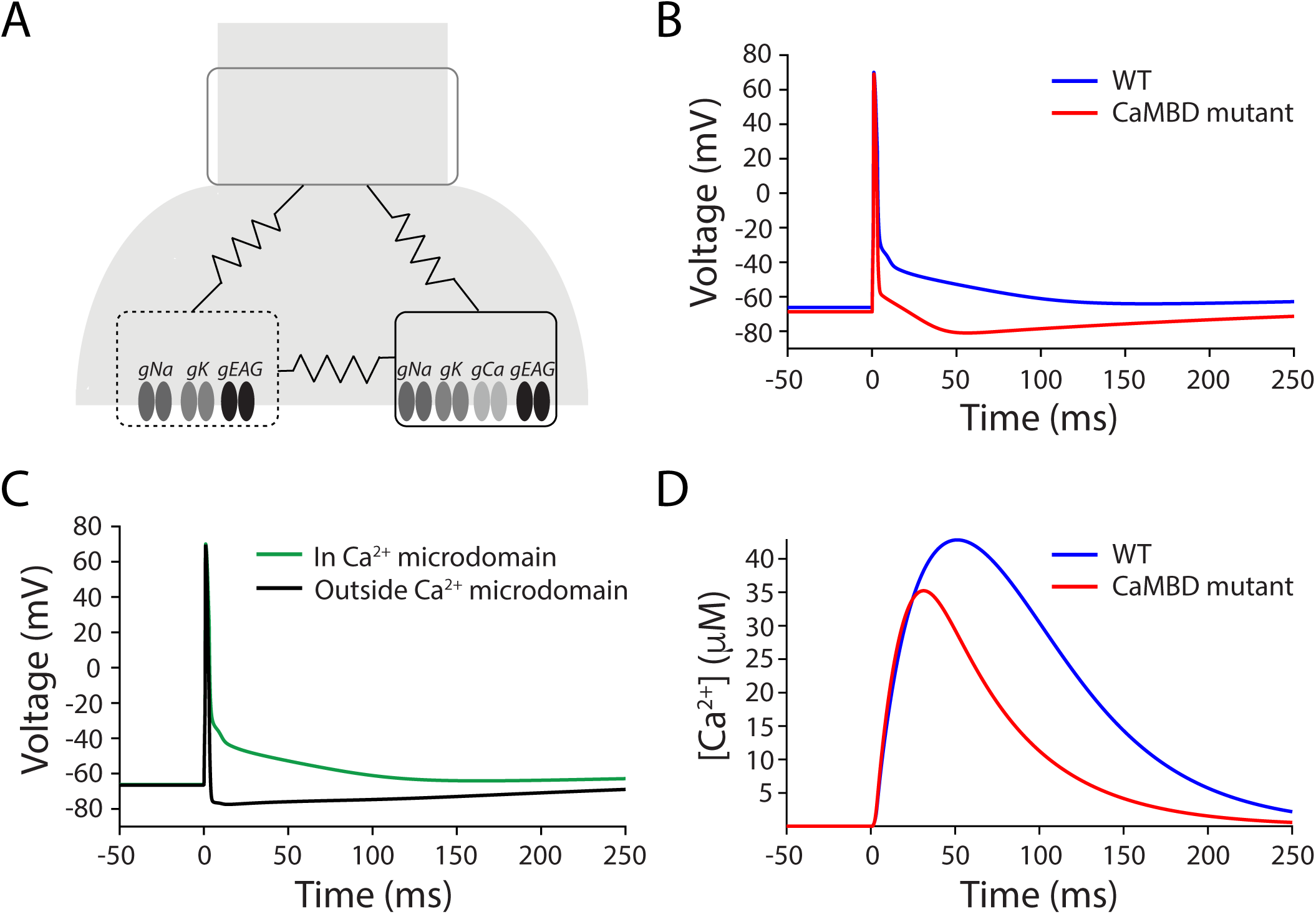
Effects of the CaMBD mutation in a three-compartment model of the presynaptic terminal. (A) Schematic of the multi-compartment model of the presynaptic terminal. The presynaptic terminal is modeled as 3 electrically coupled compartments: (1) the invading action potential (gray box, top), (2) the Ca^2+^ microdomain with voltage-gated Ca^2+^ channels and Ca^2+^ influx (black box), and (3) the parts of the synaptic terminal outside the Ca^2+^ microdomain (dotted black box). (B) Evoked potentials in response to invading action potential in the Ca^2+^ microdomain with WT EAG channels (blue) and CaMBD mutant EAG channels (red). In WT, EAG channels are inhibited by incoming Ca^2+^ and this generates a prolonged potential. The CaMBD mutant EAG channels have reduced sensitivity to Ca^2+^ and the membrane potential quickly repolarizes due to a gain of EAG function. (C) Comparison of evoked potentials in the Ca^2+^ microdomain (green) and outside the Ca^2+^ microdomain (black). Since EAG channels outside the microdomain are not inhibited by incoming Ca^2+^, they quickly repolarize the membrane after an action potential. (D) Comparison of evoked Ca^2+^ influx with WT (blue) and CaMBD mutant EAG (red). Similar to Figure 5, Ca^2+^ influx is larger and more prolonged in cells with WT EAG than in cells with CaMBD mutant EAG. All parameters of the model are identical between WT and mutant, and between the two compartments of the presynaptic terminal.

**Figure 7.**
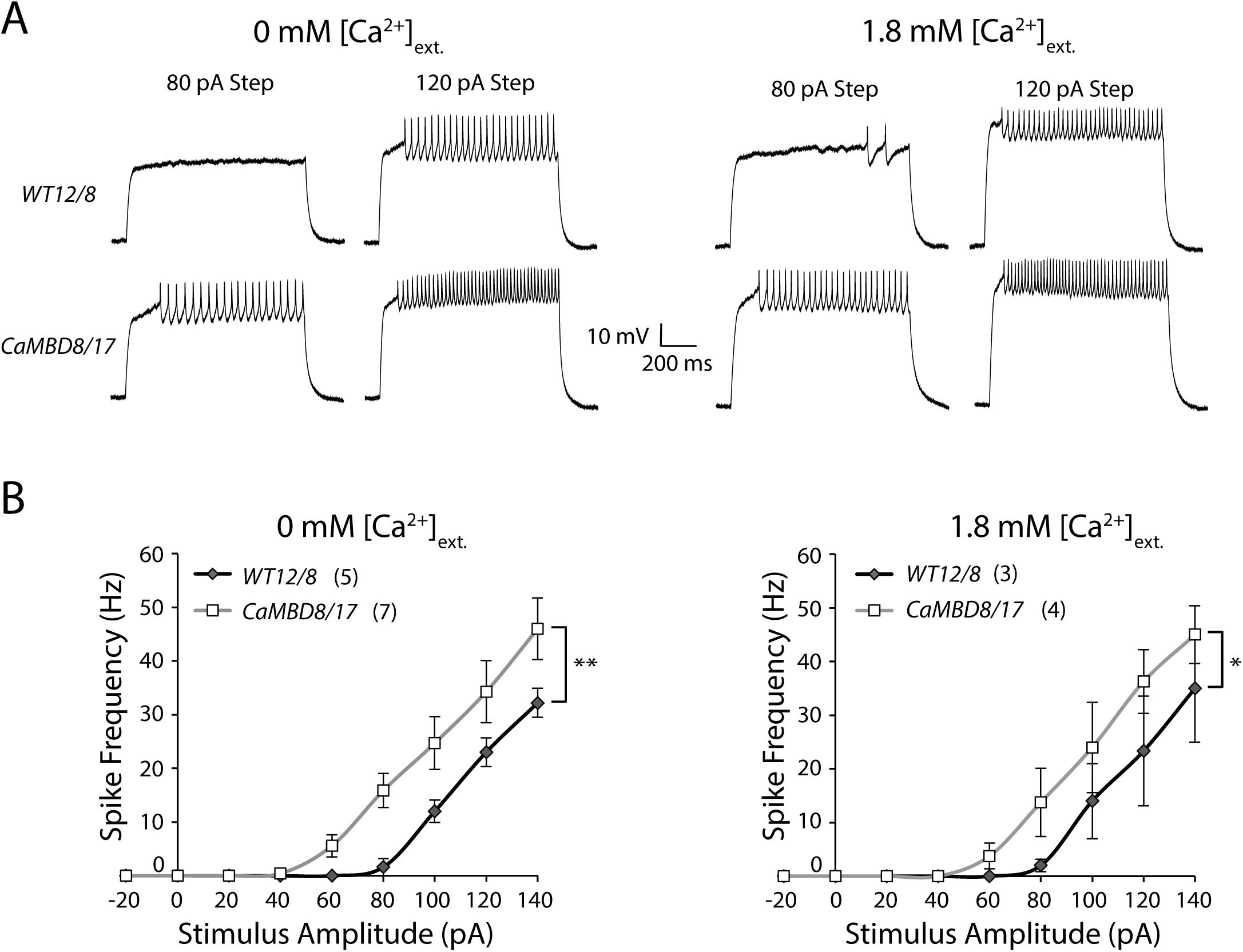
Motor neurons show somatic hyperexcitability. (A) Sample traces of whole-cell patch current clamp recordings from Type Is larval motor neuron cell bodies at selected current stimulus steps (80 pA and 120 pA) in nominally Ca^2+^-free or 1.8 mM external Ca^2+^. (B) Spike frequency vs. stimulus curves (F/I) for nominally Ca^2+^-free and 1.8 mM external Ca^2+^. There is no significant difference in the mean spike frequencies with or without Ca^2+^, p = 0.9, 3-factor ANOVA). The CaMBD mutant mean spike frequencies were significantly higher than controls (No Ca^2+^: p = 3.1 × 10^−5^; 1.8 mM Ca^2+^: p = 0.03, 2-factor ANOVA). n represents number of motor neurons recorded. Plotted values are mean ± SEM. All cells recorded in the 1.8 mM external Ca^2+^ condition were first recorded in the nominally Ca^2+^-free condition.

Recordings were performed both in nominally Ca^2+^-free external solution and in 1.8 mM external Ca^2+^. Interestingly, there was no difference with Ca^2+^ as a main effect between the two sets of F/I curves (3-way ANOVA, p = 0.9). This indicates that it is unlikely that the Ca^2+^-dependent regulation of EAG normally plays a role in somatic excitability. This could either mean that EAG is not present in the somatodendritic compartment, or that it is located too far from sites of Ca^2+^ influx to be inhibited. Given previous literature on *eag* null mutants (Srinivasan et al. 2012), it is more likely that this is an issue of differential localization.

The data from the soma present a significant contrast to those obtained at the NMJ, where there appeared to be a pronounced hypoexcitabilty. Given that these results are from different cellular compartments of the same motor neurons, it seems likely that there is compartment-specific homeostatic compensation occurring that increases drive to motor neuron terminals. The lack of effect of external Ca^2+^ suggests that the alterations in excitability do not involve acutely Ca^2+^-dependent processes. The increase in excitability may act to partially compensate for the decrease in evoked amplitude at the motor neuron terminals.

### CaMBD mutants have memory defects

The demonstration of a change in somatic excitability aimed at compensating for poor presynaptic release suggested that homeostatic changes might be able to ameliorate the effects of this mutation at the circuit level. To test this, we asked if associative memory, a complex behavior with a well-described circuitry, was disrupted in the CaMBD mutant. Adult flies were conditioned by presenting one odor in the presence of sugar, an appetitive stimulus, and a different odor with no sugar. WT flies, when asked to choose between the two odors either immediately (short-term memory) or 24 h after training (long-term memory) will choose the paired odor significantly above chance. CaMBD mutants have impairment in both types of memory (Figure 8A). Input behaviors, sugar preference and odor avoidance (Figure 8B) are not impaired. If anything, the CaMBD mutants were better than the WT control lines for avoidance of octanol. The level of short-term memory impairment was similar to that seen with *eag* null mutants, which interestingly also have enhanced octanol avoidance (C.L., data not shown). While it is not clear if the behavioral effects we see are due to the presynaptic or somatic phenotypes of this mutant, these data demonstrate that the dysregulation of excitability caused by the loss of Ca^2+^ sensitivity of the EAG channel has widespread effects on neuronal circuits and disrupts complex behaviors.

**Figure 8.**
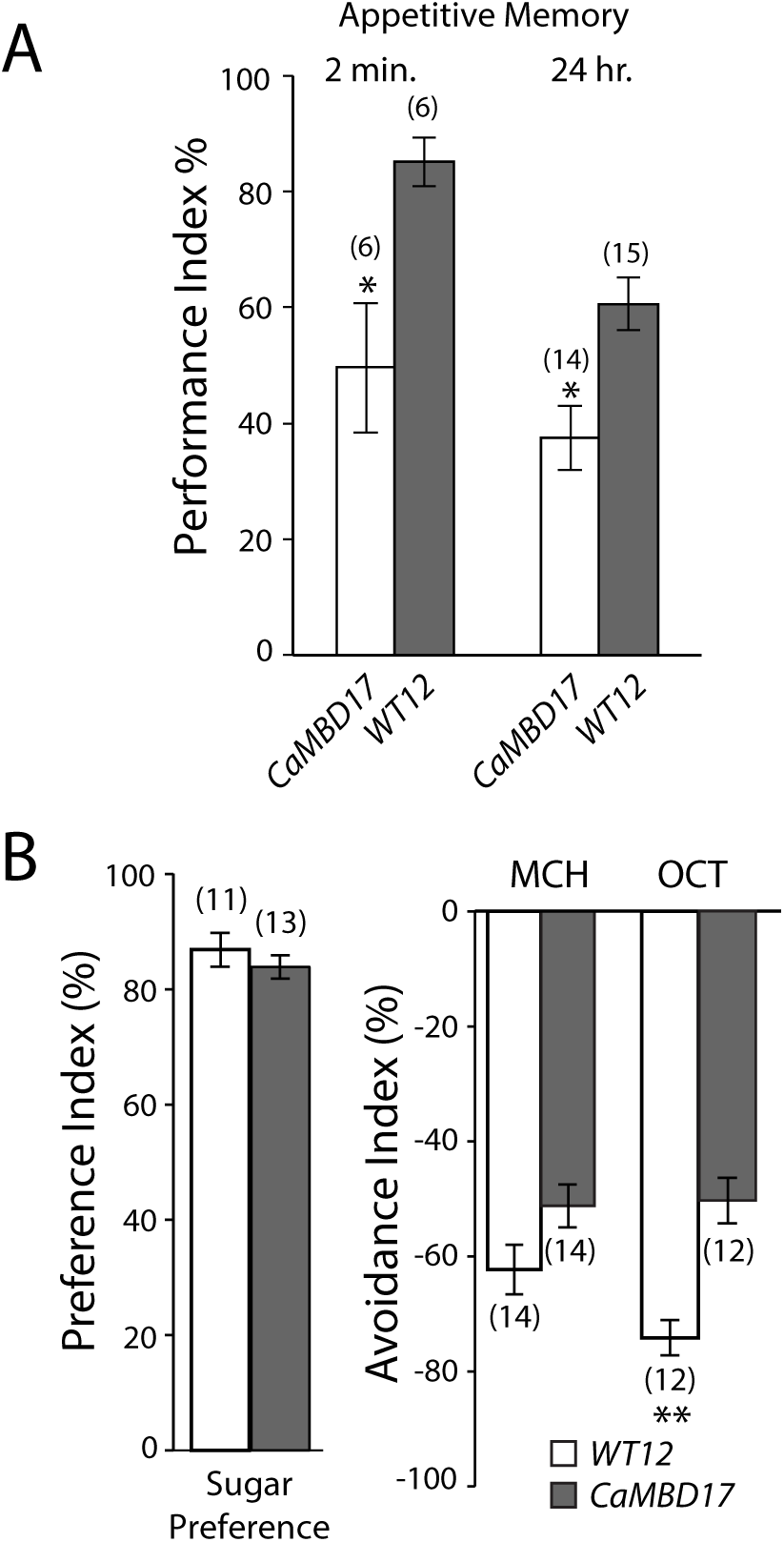
Adult behavioral phenotypes of control and CaMBD mutant adults. (A) Plot of the performance index for 2 min and 24 h memory of CaMBD mutant 17.4.1 and WT control 12.6.1 adults in a *white*^+^ genetic background. The CaMBD mutants had a significantly lower performance index for appetitive conditioning, p = 0.007 for both 2 min and 24 h tests, 2-factor ANOVA with Sidak’s multiple comparison test. n represents the number of groups tested (see methods). CaMBD mutant 8.8.1 showed a similar 2 min memory deficit, though 24 h memory was not assayed (C.L., data not shown). (B) Quantification of sugar preference (p = 0.4) and avoidance of 4-methylcyclohexanol (MCH, p = 0.08) and 3-octanol (OCT p = 0.0002). All values are mean ± SEM and comparisons done with unpaired Student’s t-test.

## Discussion

An obvious role of protein localization is to limit the action of a molecule to a particular cell compartment e.g. an anchored K^+^ channel will only affect the membrane potential of a small area surrounding it; a scaffolded kinase will preferentially phosphorylate substrates in the same complex. Less obvious consequences of binding to a scaffold are alterations of the intrinsic activities of the complexed molecules and generation of novel complex-specific effects on neuronal processes. A well-known example of this is the fusion of vesicles carrying neurotransmitters. Cytosolic bulk Ca^2+^ in neurons is typically regulated in the 10 nM to 10 μM range and only cytoplasmic Ca^2+^ microdomains close to a voltage-gated Ca^2+^ channel (VGCC) ever see Ca^2+^ at mM concentrations. The SNARE machinery has very low Ca^2+^ affinity and only functions when vesicles are docked to a complex with VGCCs (Stanley 2016). The inhibition of *Drosophila* EAG channels by Ca^2+^ is likely to be facilitated by a similar type of interaction. In this study we provide the first insights into its role by introducing mutations into the endogenous *eag* locus that block regulation by Ca^2+^.

### *Drosophila* EAG is inhibited by levels of Ca^2+^ only seen in VGCC microdomains

The fly EAG channel had been thought to be an outlier with regard to regulation by Ca^2+^. In this study we show that it is regulated in the same manner as mammalian EAG1 (Figure 1). The major difference between fly and mammalian EAGs appears to be sensitivity. Inhibition of fly EAG is only see at high μM to mM Ca^2+^ in transfected cells. It is important to note, however, that we have not measured the Ca^2+^ sensitivity *in vivo*. Although the most parsimonious expectation is that EAG Ca^2+^ sensitivity is low in *Drosophila* neurons, it remains possible that some fly-specific extrinsic factor (not present in mammalian or amphibian cells) could serve to increase it in the native context. Nonetheless, our data show a striking difference between the regulation of mammalian EAG1, which is fully inhibited at nM levels of Ca^2+^ and the fly EAG which retains activity even at mM Ca^2+^ levels. How the difference in sensitivity is achieved in the fly protein is unknown, but we present evidence that the efficacy of regulation of EAG channels by Ca^2+^/CaM is likely controlled by the C-terminal CaMBDs, as alternative splicing in the fly gene at the site of the N-terminal CaMBD does not significantly modulate apparent affinity. This Ca^2+^-dependent inhibition of EAG channels is a robust and evolutionarily conserved feature of the channel family that has so far been without demonstrated function in central neurons.

### Ca^2+^-dependent inhibition of EAG may function to tune presynaptic release

EAG channels are located in presynaptic terminals in both mammals and flies (Gillespie and Hodge 2013; Mortensen et al. 2015) and characterization of knock-outs suggest that EAG participates in repolarization after action potential invasion and controls action potential width (Wu et al. 1983). These loss-of-function studies make it clear that repolarizing K^+^ current is important for presynaptic function and that the channel has a similar overall role across species. The role of Ca^2+^-dependent inhibition of EAG currents is not as well understood. The finding that evoked release was decreased in the EAG CaMBD mutant in the face of normal spontaneous release dynamics and quantal size argued that presynaptic EAG *in vivo* is exposed to very high levels of Ca^2+^ during action potential-mediated neurotransmitter release, consistent with localization at active zones. The reduced neurotransmitter release suggests that the Ca^2+^-dependent inhibition of EAG is used to locally amplify the effects of depolarization at the active zone by decreasing the repolarizing current in that domain.

In mammals it has been speculated that the inhibition of EAG1 by Ca^2+^/CaM serves as a feedback mechanism to reduce the electrochemical driving force for Ca^2+^ influx (Schonherr et al. 2000). The idea is that EAG channels, if they were colocalized with VGCCs could control the local membrane potential after an action potential during the time that the Ca^2+^ channels are open. Inhibition of EAG by Ca^2+^ would allow the membrane potential to stay close the Ca^2+^ reversal potential and blunt the electrochemical drive for Ca^2+^ influx through the open VGCC. Our data (Figures 4 and 5) would argue against this hypothesis for the presynaptic terminal at the larval NMJ since loss of regulation by Ca^2+^ decreases presynaptic release and evoked Ca^2+^ influx. Whether this might be a viable mechanism for regulation of Ca^2+^ in dendrites would require further investigation.

Another possible role for Ca^2+^-dependent inhibition might be maintaining neurotransmitter release during repetitive firing-essentially providing a way to turn down local repolarization to maintain the ability to catalyze vesicle fusion. Our data are consistent with this type of role. Future studies of the dynamics of EAG inhibition and its dependence of action potential pattern might provide clues to this aspect of EAG function.

### Compartment-specific compensation?

The fact that presynaptic release at the NMJ and Ca^2+^ responses in central axons are reduced is consistent with the gain-of-function nature of the CaMBD mutation which allows the EAG K^+^ current to be maintained at during neuronal activation. This suggests that there is little local homeostatic change in axon terminals when the ability of Ca^2+^ to inhibit EAG currents is lost. This lack of compensation is especially interesting in light of the ability of the NMJ to quickly compensate for toxin-mediated decreases in EJP amplitude in *eag* null mutants (Bergquist et al. 2010) and to chronically compensate for EJP reductions e.g. in glutamate receptor mutants (Haghighi et al. 2003) or when muscle excitability is dampened (Paradis et al. 2001). One possible interpretation of this is that EAG itself is part of the homeostatic machinery that would be engaged in this type of neuronal insult and that its regulation by Ca^2+^ is required for compensation. Perhaps consistent with this, we saw a significant transcriptional induction of *eag* mRNA in these mutants (P. Bronk, data not shown). Another possibility is that the increase in EAG current itself occludes the effects of the normal homeostatic compensation.

While the phenotype of the nerve terminal appears relatively straightforward, somatic recordings demonstrated a surprising increase in excitability. This enhanced excitability is not directly reconcilable with the nature of the *eag* CaMBD mutation itself, since it would be expected to result in suppression of excitability due to enhanced K^+^ current. There are at least three possible scenarios that might explain this observation. One is that the loss of regulation of EAG triggers a compensation in the dendritic/somatic compartment but this compensation “overshoots” i.e. the expression of excitability-promoting channels/transporters is too high. A second possibility is that the motor neuron’s somatic/dendritic increase in excitability is a homeostatic response to the decrease in release at the NMJ. Because the normal mechanisms of compensation (locally increased presynaptic release) are not engaged, the neuron may scale up its responses to inputs to allow more transmitter to be released via increased numbers of action potentials. This would imply that there are secondary tiers of compensation and that there can be interactions between homeostatic programs similar to what has been reported when neurons receive conflicting perturbations (Bergquist et al. 2010). These possibilities are speculative, but in line with the robust homeostasis seen with other manipulations of this preparation. A third possibility is that at the action potential initiation zone, increased EAG current can act to enhance fast firing by, for example, blunting Na^+^ channel inactivation.

### Why is regulation by Ca^2+^ different in mammals and insects?

The sequence conservation of both N-and C-terminal CaMBDs from insects to mammals is striking. Why would their apparent affinities be orders of magnitude apart? The difference in sensitivity might be driven by how subcellular complexes are constituted and the relative size of synaptic compartments in the two types of organisms. The Ca^2+^-sensitivity required to carry out a function depends on the localization of the effector with respect to the Ca^2+^ source and the nature of local Ca^2+^ buffering. Channels located in very close proximity to VGCCs do not require high Ca^2+^ sensitivity, since they are exposed to very high (mM) local Ca^2+^ concentrations (Tadross et al. 2013). Conversely, EAG channels that are inhibited with high affinity could be affected by Ca^2+^ distant from the mouth of the VGCC. For fly neurons whose central processes are very small, high affinity EAG inhibition might completely abolish all EAG current and render neurons hyperexcitable, much like the case for *eag* null animals. The Ca^2+^-dependent inhibition of EAG could therefore contribute to maintaining presynaptic activity with repetitive stimulation in both species, but be tuned differently due to differences in the location of the channels, buffering capacity of the neurons or the size of the cell and synapse. A better understanding of the localizations of VGCCs and EAG channels will be required to fully understand the role Ca^2+^-dependent inhibition of EAG family channels.

### Localization of ion channels to specific subcellular domains is critical for their regulation

In neurons, ion channel-containing signaling complexes are a key class of protein assemblage. The co-localization of proteins to the same complex can strongly influence the kinetics and even the energetics of their activities and produce unforeseen interactions. Most of a neuron’s important biochemistry is not well-predicted by simply considering the equilibrium state; Ca^2+^ microdomains are an excellent example of this principle. In this study, we present data demonstrating that blocking the regulation of EAG by Ca^2+^ can disrupt the ability to form memory (Figure 8). This argues that relatively subtle changes in the regulation of excitability can have profound effects. There are an increasing number of examples of proteins whose function requires their localization to a particular microdomain (Averaimo et al. 2016; Oheim et al. 2017; Sanchez-Alonso et al. 2016). To understand how neurons work we must understand how protein complexes shape signaling and the fly EAG channel will provide a useful model for this endeavor.

**Table 1. Conditions and cell properties for electrophysiological recordings**. Values are mean ± SEM. Asterisk symbol indicates the parameter was not available for all cells. In these instances, the SEM was adjusted for the reduced value of n. The motor neuron parameters are separated by Ca^2+^ condition, but the cells in the 1.8 mM Ca^2+^ condition are cells first recorded in no external Ca^2+^ that remained healthy enough to do the second recording in external Ca^2+^.

## Acknowledgements

We would like to thank Rob Reenan (Brown University) for generously sharing *eag* constructs and for his advice on techniques for homologous recombination, and Ed Dougherty for assistance with confocal imaging.

## References

Attrill H, Falls K, Goodman JL, Millburn GH, Antonazzo G, Rey AJ, Marygold SJ, and FlyBase c. FlyBase: establishing a Gene Group resource for Drosophila melanogaster. Nucleic Acids Res 44: D786–792, 2016.

Averaimo S, Assali A, Ros O, Couvet S, Zagar Y, Genescu I, Rebsam A, and Nicol X. A plasma membrane microdomain compartmentalizes ephrin-generated cAMP signals to prune developing retinal axon arbors. Nature communications 7: 12896, 2016.

Bergquist S, Dickman DK, and Davis GW. A hierarchy of cell intrinsic and target-derived homeostatic signaling. Neuron 66: 220–234, 2010.

Budnik V, Zhong Y, and Wu C-F. Morphological plasticity of motor axons in Drosophila mutants with altered excitability. J Neurosci 10: 3754–3768, 1990.

Chen TW, Wardill TJ, Sun Y, Pulver SR, Renninger SL, Baohan A, Schreiter ER, Kerr RA, Orger MB, Jayaraman V, Looger LL, Svoboda K, and Kim DS. Ultrasensitive fluorescent proteins for imaging neuronal activity. Nature 499: 295–300, 2013.

Choi JC, Park D, and Griffith LC. Electrophysiological and morphological characterization of identified motor neurons in the Drosophila third instar larva central nervous system. J Neurophysiol 91: 2353–2365, 2004.

Dayan P, and Abbott LF. Theoretical Neuroscience: Computational and mathematical modeling of neural systems. Cambridge, MA: MIT Press, 2001.

Drysdale R, Warmke J, Kreber R, and Ganetzky B. Molecular characterization of eag: a gene affecting potassium channels in Drosophila melanogaster. Genetics 127: 497–505, 1991.

Feng Y, Ueda A, and Wu CF. A modified minimal hemolymph-like solution, HL3.1, for physiological recordings at the neuromuscular junctions of normal and mutant Drosophila larvae. J Neurogenet 18: 377–402, 2004.

Ganetzky B, Robertson GA, Wilson GF, Trudeau MC, and Titus SA. The eag family of K+ channels in Drosophila and mammals. Ann N Y Acad Sci 868: 356–369., 1999.

Ganetzky B, and Wu CF. Neurogenetic analysis of potassium currents in Drosophila: synergistic effects on neuromuscular transmission in double mutants. J Neurogenet 1: 17–28, 1983.

Gillespie JM, and Hodge JJ. CASK regulates CaMKII autophosphorylation in neuronal growth, calcium signaling, and learning. Frontiers in molecular neuroscience 6: 27, 2013.

Griffith LC, Wang J, Zhong Y, Wu CF, and Greenspan RJ. Calcium/calmodulin-dependent protein kinase II and potassium channel subunit Eag similarly affect plasticity in Drosophila. Proc Natl Acad Sci USA 91: 10044–10048, 1994a.

Griffith LC, Wang J, Zhong Y, Wu CF, and Greenspan RJ. Calcium/calmodulin-dependent protein kinase II and potassium channel subunit eag similarly affect plasticity in Drosophila. Proc Natl Acad Sci U S A 91: 10044–10048, 1994b.

Gutman GA, Chandy KG, Adelman JP, Aiyar J, Bayliss DA, Clapham DE, Covarriubias M, Desir GV, Furuichi K, Ganetzky B, Garcia ML, Grissmer S, Jan LY, Karschin A, Kim D, Kuperschmidt S, Kurachi Y, Lazdunski M, Lesage F, Lester HA, McKinnon D, Nichols CG, O’Kelly I, Robbins J, Robertson GA, Rudy B, Sanguinetti M, Seino S, Stuehmer W, Tamkun MM, Vandenberg CA, Wei A, Wulff H, Wymore RS, and International Union of P. International Union of Pharmacology. XLI. Compendium of voltage-gated ion channels: potassium channels. Pharmacol Rev 55: 583–586, 2003.

Haghighi AP, McCabe BD, Fetter RD, Palmer JE, Hom S, and Goodman CS. Retrograde control of synaptic transmission by postsynaptic CaMKII at the Drosophila neuromuscular junction. Neuron 39: 255–267, 2003.

Hara Y, Koganezawa M, and Yamamoto D. The Dmca1D channel mediates Ca(2+) inward currents in Drosophila embryonic muscles. J Neurogenet 29: 117–123, 2015.

Hardie RC. Voltage-sensitive potassium channels in Drosophila photoreceptors. J Neurosci 11: 3079–3095, 1991.

Hodgkin AL, and Huxley AF. A quantitative description of membrane current and its application to conduction and excitation in nerve. J Physiol 117: 500–544, 1952.

Immonen EV, French AS, Torkkeli PH, Liu H, Vahasoyrinki M, and Frolov RV. EAG channels expressed in microvillar photoreceptors are unsuited to diurnal vision. J Physiol 595: 5465–5479, 2017.

Kaplan W, Trout WE III. The behavior of four neurological mutants of Drosophila. Genetics 61: 399–409, 1969.

Kawasaki F, Zou B, Xu X, and Ordway RW. Active zone localization of presynaptic calcium channels encoded by the cacophony locus of Drosophila. J Neurosci 24: 282–285, 2004.

Li Y, Liu X, Wu Y, Xu Z, Li H, Griffith LC, and Zhou Y. Intracellular regions of the Eag potassium channel play a critical role in generation of voltage-dependent currents. J Biol Chem 286: 1389–1399, 2011.

Mortensen LS, Schmidt H, Farsi Z, Barrantes-Freer A, Rubio ME, Ufartes R, Eilers J, Sakaba T, Stuhmer W, and Pardo LA. KV 10.1 opposes activity-dependent increase in Ca2+ influx into the presynaptic terminal of the parallel fibre-Purkinje cell synapse. J Physiol 593: 181–196, 2015.

O’Dowd DK, and Aldrich RW. Voltage-clamp analysis of sodium channels in wild-type and mutant Drosophila neurons. J Neurosci 8: 3633–3643, 1988.

Oheim M, Schmidt E, and Hirrlinger J. Local energy on demand: Are ‘spontaneous’ astrocytic Ca2+-microdomains the regulatory unit for astrocyte-neuron metabolic cooperation? Brain Res Bull 2017.

Paradis S, Sweeney ST, and Davis GW. Homeostatic control of presynaptic release is triggered by postsynaptic membrane depolarization. Neuron 30: 737–749., 2001.

Parekh AB. Ca2+ microdomains near plasma membrane Ca2+ channels: impact on cell function. J Physiol 586: 3043–3054, 2008.

Parks AL, Cook KR, Belvin M, Dompe NA, Fawcett R, Huppert K, Tan LR, Winter CG, Bogart KP, Deal JE, Deal-Herr ME, Grant D, Marcinko M, Miyazaki WY, Robertson S, Shaw KJ, Tabios M, Vysotskaia V, Zhao L, Andrade RS, Edgar KA, Howie E, Killpack K, Milash B, Norton A, Thao D, Whittaker K, Winner MA, Friedman L, Margolis J, Singer MA, Kopczynski C, Curtis D, Kaufman TC, Plowman GD, Duyk G, and Francis-Lang HL. Systematic generation of high-resolution deletion coverage of the Drosophila melanogaster genome. Nature genetics 36: 288–292, 2004.

Ramos Gomes F, Romaniello V, Sanchez A, Weber C, Narayanan P, Psol M, and Pardo LA. Alternatively Spliced Isoforms of KV10.1 Potassium Channels Modulate Channel Properties and Can Activate Cyclin-dependent Kinase in Xenopus Oocytes. J Biol Chem 290: 30351–30365, 2015.

Sanchez-Alonso JL, Bhargava A, O’Hara T, Glukhov AV, Schobesberger S, Bhogal N, Sikkel MB, Mansfield C, Korchev YE, Lyon AR, Punjabi PP, Nikolaev VO, Trayanova NA, and Gorelik J. Microdomain-Specific Modulation of L-Type Calcium Channels Leads to Triggered Ventricular Arrhythmia in Heart Failure. Circ Res 119: 944–955, 2016.

Schonherr R, Lober K, and Heinemann SH. Inhibition of human ether a go-go potassium channels by Ca(2+)/calmodulin. Embo J 19: 3263–3271., 2000.

Srinivasan S, Lance K, and Levine RB. Contribution of EAG to excitability and potassium currents in Drosophila larval motoneurons. J Neurophysiol 107: 2660–2671, 2012.

Staber CJ, Gell S, Jepson JE, and Reenan RA. Perturbing A-to-I RNA editing using genetics and homologous recombination. Methods Mol Biol 718: 41–73, 2011.

Stanley EF. The Nanophysiology of Fast Transmitter Release. Trends Neurosci 39: 183–197, 2016.

Stansfeld CE, Roper J, Ludwig J, Weseloh RM, Marsh SJ, Brown DA, and Pongs O. Elevation of intracellular calcium by muscarinic receptor activation induces a block of voltage-activated rat ether-a-go-go channels in a stably transfected cell line. Proc Natl Acad Sci U S A 93: 9910–9914, 1996.

Sun XX, Hodge JJ, Zhou Y, Nguyen M, and Griffith LC. The eag potassium channel binds and locally activates calcium/calmodulin-dependent protein kinase II. J Biol Chem 279: 10206–10214, 2004.

Tadross MR, Tsien RW, and Yue DT. Ca2+ channel nanodomains boost local Ca2+ amplitude. Proc Natl Acad Sci U S A 110: 15794–15799, 2013.

Thibault ST, Singer MA, Miyazaki WY, Milash B, Dompe NA, Singh CM, Buchholz R, Demsky M, Fawcett R, Francis-Lang HL, Ryner L, Cheung LM, Chong A, Erickson C, Fisher WW, Greer K, Hartouni SR, Howie E, Jakkula L, Joo D, Killpack K, Laufer A, Mazzotta J, Smith RD, Stevens LM, Stuber C, Tan LR, Ventura R, Woo A, Zakrajsek I, Zhao L, Chen F, Swimmer C, Kopczynski C, Duyk G, Winberg ML, and Margolis J. A complementary transposon tool kit for Drosophila melanogaster using P and piggyBac. Nature genetics 36: 283–287, 2004.

Ufartes R, Schneider T, Mortensen LS, de Juan Romero C, Hentrich K, Knoetgen H, Beilinson V, Moebius W, Tarabykin V, Alves F, Pardo LA, Rawlins JN, and Stuehmer W. Behavioural and functional characterization of Kv10.1 (Eag1) knockout mice. Human molecular genetics 22: 2247–2262, 2013.

Wang JW, Wong AM, Flores J, Vosshall LB, and Axel R. Two-photon calcium imaging reveals an odor-evoked map of activity in the fly brain. Cell 112: 271–282, 2003.

Wang Z, Wilson GF, and Griffith LC. Calcium/calmodulin-dependent protein kinase II phosphorylates and regulates the Drosophila eag potassium channel. J Biol Chem 277: 24022–24029., 2002.

Warmke JW, and Ganetzky B. A family of potassium channel genes related to eag in Drosophila and mammals. Proc Natl Acad Sci U S A 91: 3438–3442, 1994.

Whicher JR, and MacKinnon R. Structure of the voltage-gated K(+) channel Eag1 reveals an alternative voltage sensing mechanism. Science 353: 664–669, 2016.

Wu CF, Ganetzky B, Haugland FN, and Liu AX. Potassium currents in Drosophila: different components affected by mutations of two genes. Science 220: 1076–1078, 1983.

Zhong Y, and Wu CF. Alteration of four identified K+ currents in Drosophila muscle by mutations in. Science 252: 1562–1564, 1991.

Zhong Y, and Wu CF. Modulation of different K+ currents in Drosophila: a hypothetical role for the Eag subunit in multimeric K+ channels. J Neurosci 13: 4669–4679, 1993.

Ziechner U, Schonherr R, Born AK, Gavrilova-Ruch O, Glaser RW, Malesevic M, Kullertz G, and Heinemann SH. Inhibition of human ether a go-go potassium channels by Ca2+/calmodulin binding to the cytosolic N-and C-termini. The FEBS journal 273: 1074–1086, 2006.

